# Nuclear mechanostability emerges from satellite DNA condensation into chromocenters

**DOI:** 10.1101/2025.08.07.669059

**Authors:** Franziska B. Brändle, Benjamin Frühbauer, Ilaria Ceppi, Logan de Monchaux-Irons, Thor van Heesch, Alessia Sommer, Francesco Rivetti, Sung Sik Lee, Chiara Morelli, Paolo Arosio, Anna Sintsova, Thomas C.T. Michaels, Petr Cejka, Jocelyne Vreede, Madhav Jagannathan

**Affiliations:** Institute of Biochemistry, ETH Zürich; Zürich, Switzerland; Bringing Materials to Life Initiative, ETH Zürich; Zürich, Switzerland; Institute for Research in Biomedicine, Università della Svizzera italiana; Bellinzona, Switzerland; Van’t Hoff Institute for Molecular Sciences; Amsterdam, Netherlands; Scientific Center for Optical and Electron Microscopy (ScopeM); Zürich, Switzerland; Institute of Chemical and Bioengineering, ETH Zürich; Zürich, Switzerland; Institute of Microbiology, ETH Zürich; Zürich, Switzerland

**Keywords:** Satellite DNA, Chromocenters, Nuclear mechanostability, Heterochromatin, Phase separation

## Abstract

As the largest organelle, the nucleus endures significant mechanical stresses over the cellular lifespan, and mechanostability, i.e. the ability to resist deformation, is critical for genome integrity and function. Here, we reveal that nuclear mechanostability is an emergent property arising from the clustering of satellite DNA repeats into nuclear condensates known as chromocenters. Targeted chromocenter disruption in Drosophila testes subjected to natural and artificial mechanical stress compromises nuclear mechanostability, leading to deformed nuclei, DNA damage, and chromosome breaks. Conversely, enhancing chromocenter coalescence through genetic means improves mechanostability. Molecular dynamics simulations suggest that chromocenters enable physically linked chromosomes to respond collectively, rather than individually, to mechanical challenge, and dissipate external forces over a larger nuclear surface. We propose that the satellite DNA-dependent mechanostability framework described here likely extends to other cells and tissues facing mechanical stress, and offers an explanation for the evolutionary success of these non-coding repeats across eukaryotes.

## INTRODUCTION

As the largest cellular organelle, the nucleus experiences substantial mechanical forces from confinement and compression during migration, as well as stretching and shear stress within tissue environments ^1,2^. In these contexts, nuclear mechanostability, i.e. the ability of nuclei to resist mechanical deformation, protects the genetic information from damage^1–5^. Loss of nuclear mechanostability leads to increased DNA damage, contributing to senescence in non-transformed cells and invasiveness in cancer cells^6,7^. The mesh-like nuclear lamina at the inner nuclear membrane and the chromatin polymer within the nucleus are considered to be major factors governing nuclear mechanostability^8–13^. Generally, the lamina meshwork, consisting of A-type and B-type lamins, is thought to resist large deformations^8,13^. Of these, the A-type lamins play the dominant role in nuclear mechanostability and their dysfunction leads to deformed nuclei in many cells and tissues and is associated with pathological outcomes^9,12,14–19^. However, A-type lamins are expressed at very low levels in undifferentiated cells^20,21^ suggesting alternative means to maintain nuclear mechanostability. Notably, the spring-like chromatin, especially at constitutive heterochromatin, has been implicated as a lamina-independent mechanism to protect the nucleus from mechanical stress^8,13,22–26^. Accordingly, local chromatin compaction by heterochromatin-associated histone modifications, heterochromatin cross-linking by heterochromatin protein 1 (HP1), HP1 condensation and peripheral heterochromatin tethering are all thought to contribute to nuclear mechanostability^8,22,25–28^.

On a more global scale, the organization of constitutive heterochromatin varies between cell types and species, ranging from ‘wetting’ the surfaces of the nuclear envelope and nucleolus (e.g. cultured human cells) to coalescence into compact DAPI-dense nuclear foci known as chromocenters (e.g. *Drosophila*) (Fig. 1A). These context-specific differences in higher-order constitutive heterochromatin organization are likely dependent on the extent of heterochromatin tethering to the nuclear envelope through factors such as Lamin A, the Lamin B receptor (LBR) and the Lamina-associated Polypeptide 2 beta (LAP2β)^29–32^. Indeed, processive elimination of heterochromatin tethers in human cells results in the formation of chromocenter-like DAPI-dense foci^32^. In addition, the degree of chromocenter coalescence, i.e. the number of chromocenters per nucleus, also varies between cells of the same organism, with more differentiated cells generally exhibiting increased coalescence into fewer chromocenters^33,34^. However, the phenotypic effects of these differences in higher order heterochromatin organization have not been fully investigated.

**Figure 1.**
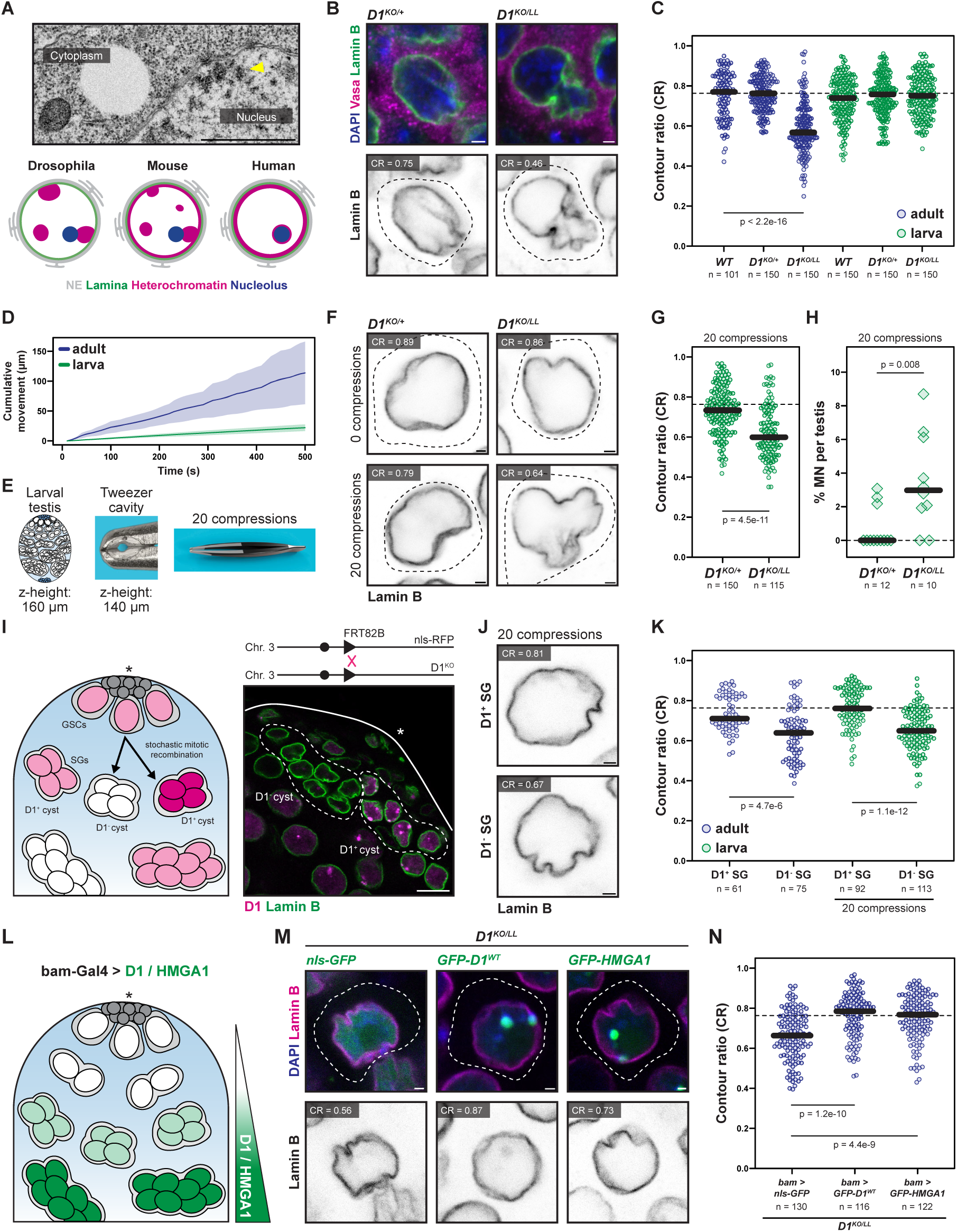
The satellite DNA-binding protein D1 promotes nuclear mechanostability in *Drosophila* male germ cells. (A) Top: Transmission electron microscopy (TEM) image of a section of a *Drosophila* nucleus from spermatogonia-enriched testes (*nos > bam^RNAi^*). Arrowhead points to a heterochromatin cluster at the nuclear periphery. Bottom: Schematic of the continuum of constitutive heterochromatin organization in *Drosophila*, mouse and human cells with the nuclear envelope (gray), the lamina (green), heterochromatin (magenta) and the nucleolus (blue) depicted. (B) Spermatogonia from control (*D1^KO/+^*) and *D1* mutant (*D1^KO/LL^*) adult testes stained for Lamin B (green), Vasa (magenta) and DAPI (blue). Outlines indicate cell boundaries. (C) CR for spermatogonia from the indicated genotypes and developmental stages. (D) Cumulative movement of larval versus adult testes over time. (E) Schematic of the tweezers used to compress larval testes. (F) Single spermatogonia from control and *D1* mutant larval testes with and without 20 compressions stained for Lamin B. Outlines indicate cell boundaries. (G, H) Spermatogonia CR (G) and % MN per testis (H) from the indicated genotypes after 20 compressions. (I) Schematic (left) and example (right) of D1-positive and D1-negative spermatogonia within the same tissue generated using FLP-FRT-based mitotic recombination. The adult testis (right) was stained for D1 (magenta) and Lamin B (green) with the outlines indicating D1-positive and D1-negative spermatogonial cysts. (J) D1-positive and D1-negative larval spermatogonia post compression and stained for Lamin B. (K) CR for D1-positive and D1-negative spermatogonia in adult and compressed larval testes. (L) Schematic of rescue experiment in an adult *D1^KO/LL^* background. (M) Adult *D1* mutant spermatogonia expressing NLS-GFP/GFP-D1^WT^/GFP-HMGA1 (green) co-stained for Lamin B (magenta) and DAPI (blue). (N) CR quantifications of the indicated rescue constructs. All scale-bars are 1 µm (except panel I, 10 µm). Indicated p values are from Mann-Whitney U-tests. For the CR and MN plots, the thick black line indicates the median while the dashed line represents the median CR (0.76) in adult *D1* heterozygous spermatogonia.

Unlike other features of constitutive heterochromatin organization, the contribution of chromocenters to nuclear mechanostability remains poorly understood, since these foci are remarkably hard to manipulate and generally resistant to perturbations such as loss of constitutive heterochromatin-associated histone modifications and heterochromatin-associated proteins^22,23,35,36^. Although reduced chromocenter compaction has been recently linked to nuclear mechanics in cultured mouse cells^37^, these experiments relied on chemical inhibition of H3K9 trimethylation, making it hard to distinguish chromocenter-specific contributions from other heterochromatin-related functions. Moreover, we still do not understand whether the degree of chromocenter coalescence affects the ability of nuclei to withstand mechanical forces. Finally, whether chromocenters influence nuclear mechanostability in a native organismal setting also remains unresolved.

One of the main constituents of constitutive heterochromatin and chromocenters are non-coding tandem repeats known as ‘satellite DNA’^33^. In many species including plants, insects and mammals, satellite DNA repeats from two or more chromosomes are clustered into individual chromocenters, which exhibit condensate-like behaviour depending on the cell type^35,38–40^. Notably, sequence-specific satellite DNA-binding proteins have been identified recently as factors that are required for chromocenter formation in *Drosophila* and mouse cells^41,42^. In *Drosophila*, two such proteins, D1 and Prod, have been shown to cluster their cognate satellite DNA repeats. Intriguingly, chromocenters in most tissues are unaffected by the loss of either D1 or Prod^42^, suggesting functional redundancy between these two satellite DNA-binding proteins. However, *D1* mutation alone is sufficient to disrupt chromocenters in early male germ cells, hereafter spermatogonia ^41,42^. Thus, modulating D1-dependent satellite DNA clustering provides a unique opportunity to determine chromocenter function in a native setting.

In this study, we demonstrate that the condensation of satellite DNA into chromocenters safeguards nuclear mechanostability and protects genome integrity in mechanically challenged *Drosophila* cells. Using molecular dynamics (MD) simulations, *in vitro* assays using purified proteins and *in vivo* mutational scanning, we show that complex coacervation, a charge-driven phase separation process, underlies D1-dependent chromocenter formation. We use these newly generated insights to selectively tune both chromocenter compaction and coalescence *in vivo*. In a tissue facing mechanical stress, we reveal that reducing the compaction and coalescence of chromocenters renders cells vulnerable to both endogenous and exogenous forces, with these cells exhibiting severe nuclear deformations, chromosome breakage and loss of genome integrity. In contrast, enhancing the coalescence of chromocenters protects vulnerable nuclei from deformation. We propose that chromocenter-dependent nuclear mechanostability emerges from satellite DNA-dependent physical connections between chromosomes and likely synergizes with HP1-dependent chromatin cross-links, peripheral heterochromatin tethering and overall heterochromatin levels to protect genome integrity in mechanically challenged cells and tissues.

## RESULTS

### Chromocenter disruption leads to mechanical stress-induced nuclear deformation

*Drosophila* male germ cells known as spermatogonia require the satellite DNA-binding protein D1 to cluster the abundant AATAT satellite DNA repeat into chromocenters (Fig. S1A)^41^. Here, we used a newly generated knockout allele of *D1* (*D1^KO^*) in combination with a previously validated mutant allele (*D1^LL03310^*) to assess chromocenters in *D1* mutant (*D1^KO/LL^*) spermatogonia (Fig. S1B). Loss of D1 resulted in increased number of AATAT foci per cell (declustering, Fig. S1C, D). These foci were also dispersed over a larger volume of the nucleus (decompaction, Fig. S1E). Interestingly, we observed that a large fraction of *D1* mutant spermatogonia displayed severe nuclear deformations (Supplementary Video 1). To quantify this phenotype, we employed a metric termed the ‘nuclear contour ratio’ (CR)^15,43^, also referred to as nuclear circularity or nuclear form factor^44^, which is calculated as 4π x area/perimeter^2^. A CR value of 1.0 represents a perfect circle, while CR values below 1.0 reflect cells with increasingly deformed nuclei. Spermatogonia from control (*D1^KO/+^*) adult testes exhibited a median CR of 0.76, which is comparable to wild-type adult spermatogonia (median CR = 0.77), and reflects the natural level of nuclear deformation observed in these cells (Fig. 1B, C). In contrast, *D1* mutant spermatogonia exhibited a median CR of 0.57, a significant change in comparison to the controls and consistent with the severe deformations observed in this sample (Fig. 1B, C). Consistent with previous reports^41^, we observed that 10% of D1 mutant spermatogonia contained micronuclei (Fig. S1F).

Altered membrane lipid composition^45^, compromised mechanostability^15,18,46^, osmotic shock^47^ and chromosome mis-segregation^48^ are all known to cause nuclear deformations. In *Drosophila*, a contractile muscle sheath forms around the testis during metamorphosis, imparting peristaltic contractions that facilitate the movement of sperm through the testis tubule. Consistently, live imaging revealed recurrent contractions in adult testes, while larval testes, which are embedded in a fat body, exhibited little to no movement over the same period (Fig. 1D, Supplementary Video 2). In addition, we observed that both control and *D1* mutant larval spermatogonia exhibited wild-type nuclear morphology and did not contain micronuclei (Fig. 1C, Fig. S1F), even though D1 protein levels were effectively depleted in *D1* mutant larval testes (Fig. S1B). These data suggested that the nuclear deformations observed in adult *D1* mutant spermatogonia may arise due to mechanical challenges imposed by the surrounding muscle sheath.

We therefore set out to determine if there were causal relationships between chromocenter disruption in *D1* mutant spermatogonia, nuclear deformation in the same cells and the mechanical stress experienced by the adult testis. To do so, we established a method to impart mechanical stress on demand and in a controlled manner on larval testes. Specifically, we developed specialized tweezers with a central cavity of 140 µm, which is ∼20 µm smaller than the average height of larval testes (Fig. 1E). We used these tweezers to compress control and *D1* mutant larval testes 20 times in ∼15-20 seconds, to mimic the mechanical forces imposed by the muscle sheath in the adult tissue. The time lag between the compressions and fixation was, at most, 5 minutes. Strikingly, these tweezer-mediated compressions induced nuclear deformations specifically in the *D1* mutant larval spermatogonia (Fig. 1F, G), with these deformations reliably observed across individual testes (Fig. S1G). These mechanical compressions were also sufficient to induce micronuclei formation in larval spermatogonia lacking D1 (Fig. 1H). To ensure that the nuclear deformation in *D1* mutant larval spermatogonia was independent of other confounding factors such as testis size variation or uneven force application, we generated mosaic testes containing D1-positive and D1-negative cells within the same tissue using the FLP-FRT system (Fig. 1I, Fig. S1H). Importantly, we found that D1-negative spermatogonia exhibited increased deformations in both adult testes and compressed larval testes, in comparison to their D1-positive counterparts (Fig. 1J, K). Thus, the deformations observed in D1 mutant spermatogonia can be induced through the application of mechanical force, indicating that D1 is important to safeguard nuclear mechanostability.

We next sought to determine whether it is chromocenter formation or an alternative function that underlies D1-dependent nuclear mechanostability. To address this, we first expressed GFP-tagged D1^WT^ in mid-late adult spermatogonia in a *D1* mutant (*D1^KO/LL^*) background (Fig. 1L-N). Importantly, GFP-D1^WT^ expression fully restored spermatogonial nuclear morphology (Fig. 1M, N) in this background. Next, we turned to HMGA1, a mammalian DNA-binding protein that clusters satellite DNA repeats into chromocenters ^41,49^ but only shares ∼11% sequence identity with D1. Despite hundreds of millions of years of evolution separating mammals and *Drosophila*, we remarkably found that GFP-tagged HMGA1 formed D1-like DAPI-dense foci when expressed in *D1* mutant adult spermatogonia (Fig. 1M, Fig. S2A-C). Moreover, GFP-HMGA1 expression was sufficient to rescue nuclear deformations in the absence of D1 (Fig. 1M, N). These data specifically suggest that satellite DNA clustering into chromocenters is the main component of D1-dependent nuclear mechanostability *in vivo*.

### Mechanically induced nuclear deformation is associated with DNA damage and chromosome fragmentation

We then assessed the stability of chromocenters in larval spermatogonia under mechanical stress. We found that AATAT clustering was largely intact in *D1* mutant larval spermatogonia in the absence of tweezer-induced compressions (Fig. S3A, B). However, the clustering of AATAT was disrupted in the same cells post compression, suggesting that D1 maintains chromocenter stability in response to mechanical stress (Fig. S3A, B). In adult testes, which experience persistent mechanical stress due to the contractile muscle sheath, we observed elevated levels of DNA damage (γH2Av foci) in *D1* mutant spermatogonia (Fig. S3C, D). We also assessed what was contained within micronuclei by labeling centromeres (CenpC-tdTomato) and telomeres (HOAP-GFP). While 78.9% of micronuclei contained telomeric foci, only 39.6% contained centromeric foci, suggesting that many micronuclei contain chromosome fragments (Fig. S3E, F). FISH against chromosome-specific satellite DNA repeats also revealed that micronuclei formation was not chromosome-specific (Fig. S3F). We suggest that in the absence of D1, the chromosomes contained within a mechanically compromised nucleus are prone to breakage, and the resultant fragments are susceptible to blebbing out from the interphase nucleus. While the precise events underlying the DNA damage are not yet completely clear, these observations reinforce the notion that D1-dependent chromocenter formation is critical for nuclear mechanostability and genome integrity.

### The LINC complex mediates nuclear deformation in D1 mutant spermatogonia

In eukaryotes, force transmission across the nuclear envelope relies on the <u>li</u>nker of the <u>n</u>ucleoskeleton and <u>c</u>ytoskeleton (LINC) complex, which consists of membrane-anchored SUN and KASH proteins^10,50,51^. In cells with compromised nuclear mechanostability, the activity of the LINC complex results in cytoskeleton-mediated nuclear deformations^9,14^. We found that loss of the SUN domain gene *klaroid* (using a validated deletion allele, *koi*^80^)^52^ did not cause a significant change in spermatogonial nuclear morphology (Fig. 2A, B). However, we observed that the loss of *koi* in *D1* mutant spermatogonia largely abolished nuclear deformations (Fig. 2A, B). In addition, we observed fewer micronuclei in the *koi* and *D1* double mutant spermatogonia (Fig. 2C). These data suggest that the deformations in *D1* mutant nuclei occur in response to actin filament- or microtubule-dependent cytoskeletal forces that are transduced by the LINC complex.

**Figure 2.**
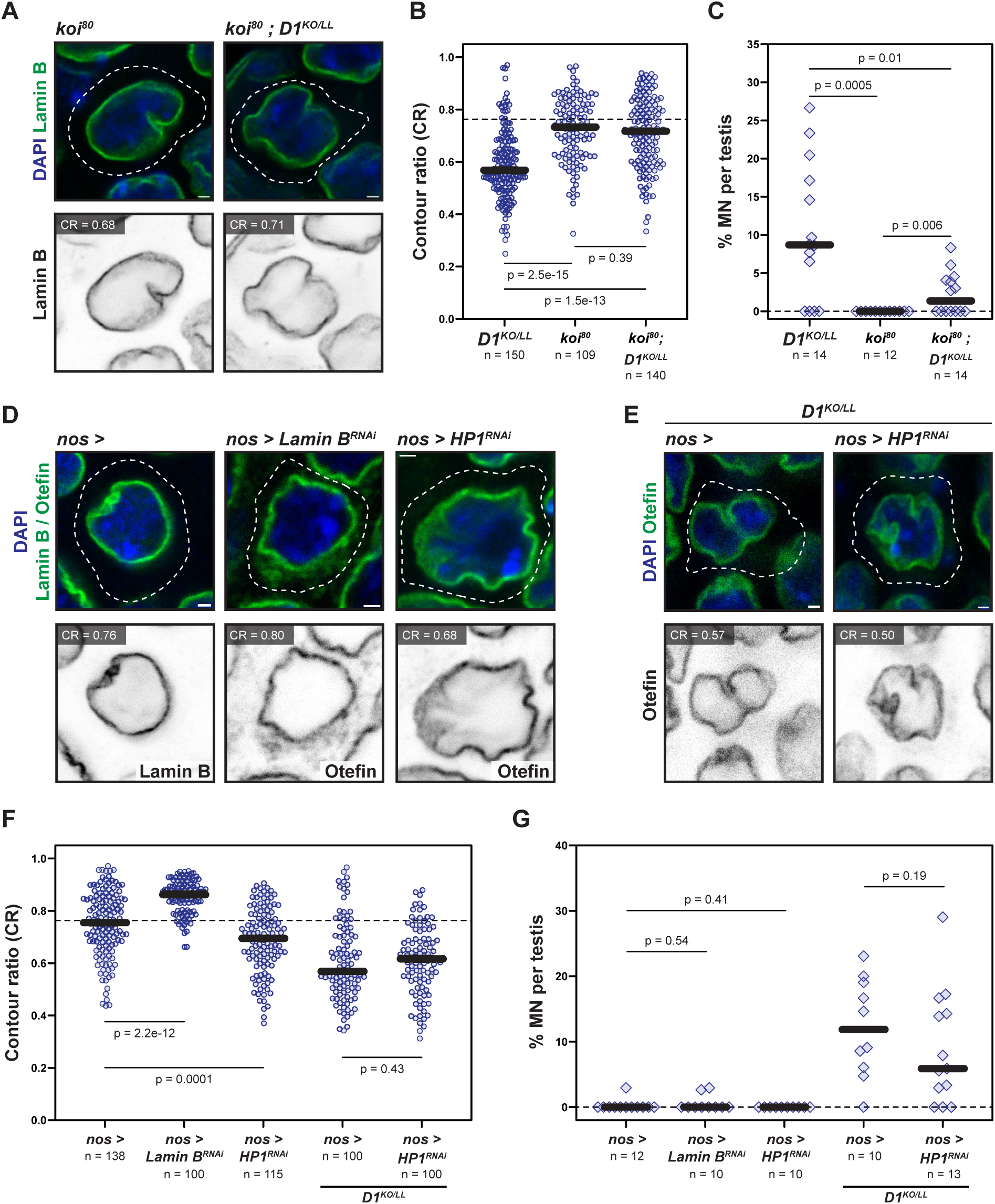
The interplay between D1 and known nuclear mechanostability factors. (A) Adult spermatogonia from the indicated genotypes stained for Lamin B (green) and DAPI (blue). (B, C) CR of adult spermatogonia (B) and % MN per testis (C) from the indicated genotypes. *D1^KO/LL^* data replotted from Fig. 1C and Fig. S1F. (D, E) Adult spermatogonia from the indicated genotypes were stained for Lamin B/Otefin (green) and DAPI (blue). (F, G) CR of adult spermatogonia (F) and % MN per testis (G) from the indicated genotypes. All scale-bars are 1 µm. Indicated p values are from Mann-Whitney U-tests. For the CR and MN plots, the thick black line indicates the median while the dashed line represents the median CR (0.76) in adult *D1* heterozygous spermatogonia.

### D1 promotes nuclear mechanostability independently of the nuclear lamina

The nuclear lamina and HP1 are known factors that contribute to nuclear mechanostability^15,19,25,27^. Since undifferentiated spermatogonia do not express the A-type lamin at significant levels, we first assessed nuclear deformation following depletion of Lamin B (Fig. 2D, Fig. S4A, C). Surprisingly, we found that Lamin B depletion, in an otherwise WT background, did not have an adverse effect on spermatogonial nuclear morphology (Fig. 2F). In addition, Lamin B depletion did not lead to a significant occurrence of micronuclei in spermatogonia (Fig. 2G). These findings echo recent observations in mammalian cells that acute Lamin depletion does not have a substantial effect on nuclear integrity, in part due to redundancy with other NE-associated proteins^53^. Finally, overexpression of GFP-tagged Lamin B did not rescue nuclear deformations in *D1* mutant spermatogonia (Fig. S4D). These data suggest that Lamin B has a negligible effect on spermatogonial nuclear morphology.

In contrast, we found that depletion of HP1 (Fig. 2D, Fig. S4B, C) led to a small but significant increase in nuclear deformation (Fig. 2F, median CR = 0.70). However, micronuclei were not observed following HP1 depletion (Fig. 2G) and we also noted that the number of D1 foci in spermatogonia was not altered (Fig. S4E). These data suggest that HP1 depletion does not affect chromocenter formation, which is consistent with previous reports^35,36^, and highlight that the effect of HP1 depletion on spermatogonial nuclear morphology is more modest than that of loss of *D1*. Consistently, spermatogonia lacking both HP1 and D1 exhibited nuclear deformation and micronuclei to a similar extent to what we observed in *D1* mutant spermatogonia alone (Fig. 2E-G).

Since satellite DNA repeats and chromocenters have been implicated in the regulation of gene expression^54,55^, we tested whether loss of D1 altered expression of Lamin B or HP1 (encoded by *Su(var)205* in *Drosophila*). Specifically, we performed RNA sequencing on spermatogonia-enriched (*nos > bam^RNAi^*) adult testes in the presence and absence of D1 (Fig. S4F). We noted that the proportion of spermatogonia was decreased in *D1* mutant testes in comparison to the control (Fig. S4F), likely due to known germ cell viability defects in the absence of D1^41^. To minimize the effect of these tissue composition differences, we limited our differential expression analyses to genes that passed a relatively stringent cutoff (log_2_FC>2, p_adj_<0.05) (Fig. S4G). In total, we only observed modest changes in gene expression in the absence of D1 – 321 upregulated and 101 downregulated genes. Although *Lamin B* and *Su(var)205* were not differentially expressed based on our cutoff, we noticed that the expression of both genes was subtly decreased in the *D1* mutant in comparison to the control (Fig. S4G). While these subtle transcriptional changes likely stem from tissue composition differences (Fig. S4F), we nevertheless assessed whether there was a corresponding reduction in protein level. Specifically, we quantified Lamin B and HP1 protein levels in mosaic testes containing D1-positive and D1-negative spermatogonia. Our data clearly indicate that the absence of D1 does not alter the protein levels of Lamin B or HP1 in spermatogonia (Fig. S4H). Taken together, our data suggest that D1-dependent chromocenter formation promotes nuclear mechanostability in spermatogonia independently of Lamin B and more effectively than HP1.

### Coarse grained polymer modeling reveals how chromocenters impact nuclear mechanostability

Coarse grained polymer models have been successfully used to illuminate how chromatin, especially features such as heterochromatin levels, peripheral tethering and cross-linking, functions in nuclear mechanostability^25,56,57^. Therefore, to obtain an intuition on how chromocenters influence nuclear mechanostability, we turned to a polymer modeling approach and adapted a previously described coarse-grained model of the nucleus^58^. In our model, we took a reductionist approach and intentionally excluded factors such as peripheral tethering, cross-links and heterochromatin types to distinguish the role of chromocenters from other well-known roles of heterochromatin in nuclear mechanics. Specifically, we modeled chromosomes as beads-on-a-string, with each chromosome consisting of 1000 monomers and with the first 100 monomers of each chromosome simply designated as ‘satellite DNA’ (Fig. 3A). These chromosomes were encapsulated within an elastic polymeric shell, mimicking the nuclear envelope/lamina (Fig. 3B). We used 20 chromosomes for our modeling to increase the dynamic range of the number of chromocenters per nucleus. These chromosomes occupied 25% of the shell (nuclear) volume, which is in line with measurements made in intact *Drosophila* nuclei^59^.

**Figure 3.**
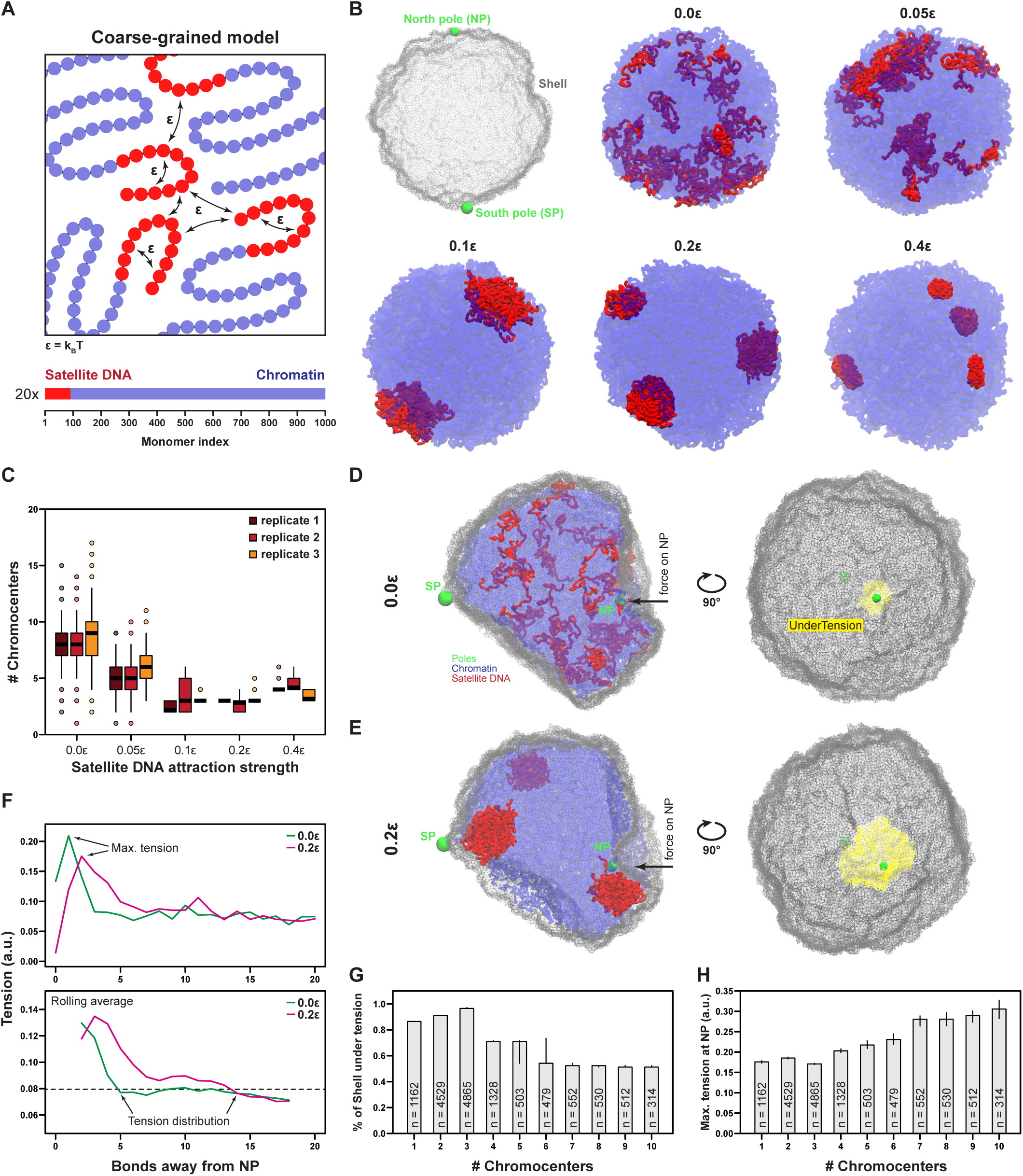
Coarse-grained polymer modeling reveals a role for chromocenter-like clustering in the dissipation of mechanical force. (A) Schematic of the simulated chromosomes. (B) Representation of the polymeric shell (top left) and chromatin organization at different levels of satellite DNA attraction strength after 7.5e+7 simulation steps (ɛ = k_B_T). NP and SP refer to diametrically opposite points on the shell. (C) Quantification of chromocenter-like structures observed in three independent equilibrium simulations with varying satellite DNA attraction strengths. (D, E) Visualization of ‘compressed’ nucleus in the absence (D) and presence (E) of chromocenter-like clustering. Shown are a cross-section (left) and a top view of the membrane (right). The yellow surface (right) highlights the shell area under increased bond tension. (F) Bond tension at the encapsulating shell from (E) as a function of distance to the NP in the absence (green) and presence (magenta) of chromocenter-like structures. Top: maximum tension around the NP. Bottom: Rolling average of bond tension with a window size of 5, revealing the return of the tension to the cutoff value (dashed line). (G, H) Tension dissipation away from the NP (G) and maximum tension at the NP (H) as a function of chromocenter-like clustering. Error bars show the 95% confidence intervals.

We simulated chromocenter-like structures, defined as satellite DNA from different chromosomes within a distance cutoff (see Materials and Methods), by tuning the attraction (ε) between satellite DNA monomers (Fig. 3A). Even in the absence of attraction (ε=0), the lowest number of chromocenters was generally 8-10, likely due to the random encounters of satellite DNA monomers from multiple chromosomes (Fig. 3B, C). However, we observed chromocenters that resembled their cellular counterparts when we increased the attraction strength (ε=0.05 – 0.4) (Fig. 3B, C). Interestingly, the number of chromocenters is lowest at intermediate attraction strengths (ε = 0.1 – 0.2), likely because higher attraction strengths favor satellite DNA interactions within the same chromosome (Fig. 3B, C). We also found that the chromocenters in our polymer model were often positioned at the shell periphery, consistent with observations made in cells^34^. This peripheral positioning may occur due to ‘frustration’ of phase separation by polymer networks e.g. chromatin^60,61^. At the same time, we note that these peripheral chromocenters are not physically tethered to the encapsulating shell, a feature recently shown to be important for the contribution of heterochromatin to nuclear mechanics^57^.

To simulate mechanical challenge, we designated opposite poles of the encapsulating shell as north pole (NP) and south pole (SP) (Fig. 3B). The SP was immobilized by a wall potential tangent to the shell surface and an iteratively increasing force was applied on the NP towards the SP, 20 times, with pauses between the force applications. To estimate the strain exerted by the applied force on the shell, we measured bond-length deviations from equilibrium values of connected shell monomers, hereafter referred to as tension. In the absence of satellite DNA clustering, we found that the tension on the shell was high and largely confined to the immediate vicinity of the north pole (Fig. 3D, F). In contrast, the formation of chromocenters led to the tension exerted by the same force being dissipated across a larger area of the shell (Fig. 3D-F), along with a reduction in the maximum tension experienced at the NP (Fig. 3F). Notably, this force dissipation scaled with increased satellite DNA clustering and a corresponding reduction in chromocenter number (Fig. 3G, H). The displacement of spherical shells under a point load has been previously described theoretically^62,63^ and relates the length scale of this dissipation, *l_p_*, to the shell’s elastic modulus *E*, radius *R*, thickness *h* and its internal pressure *p*,

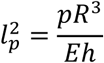

Given that the material properties and dimensions of the shell do not change with increased satellite DNA clustering, we suggest that the increase in force dissipation arises due to elevated internal pressure due to changes in the chromatin organization upon chromocenter formation. Overall, our simulations support the notion that chromocenters facilitate the dissipation of mechanical force and likely function along with other features of heterochromatin organization to preserve nuclear mechanostability. Importantly, the modeling predicts that the degree of chromocenter coalescence should scale positively with nuclear mechanostability.

### D1 phase separates in vitro through a complex coacervation-like process

In order to test the predictions of the polymer modelling data (Fig. 3G, H), we set out to obtain a molecular understanding of D1-dependent chromocenter formation, with the goal of modulating chromocenter coalescence in intact tissues. Although predicted to be fully disordered (Fig. 4A, B), D1 contains 11 DNA-binding AT-hook motifs that are distributed throughout the protein (Fig. 4B, C)^64^. To understand how D1 engages with DNA, we used atomistic molecular dynamics (MD) simulations to determine the bound conformations of all 11 AT-hooks on 3xAATAT DNA. Specifically, we conducted our simulations using the Amber99sb-ildn bsc0 cufix force field^65,66^, which performed optimally in simultaneously recapitulating DNA structure, protein disorder, and protein-DNA interactions (Fig. S5A-I). Importantly, the predicted ensembles mirrored existing structures of single AT-hooks bound to DNA (Fig. S5E-I, Fig. S6), highlighting that this method captures the relevant features of this interaction. Subsequently, we estimated the relative binding strengths of each of the AT-hooks using a recently described steered MD simulation protocol^67^ to estimate the free energy of dissociation of the AT-hooks from the DNA strand (Fig. S7A-F). Our results suggest that D1 contains AT-hooks with an ∼3-fold range in DNA-binding affinities (Fig. 4C), likely influenced by the sequence composition around the core Gly-Arg-Pro (GRP) motif. Interestingly, we noted that groups of D1 AT-hooks were separated by regions consisting of negatively charged residues (Fig. 4B). To determine the effect of these negatively charged regions (NCRs) on D1 binding to DNA, we first simulated DNA-binding for a peptide consisting of two AT-hooks (#3 and #4) separated by the 24 residue NCR1 (Fig. 4D). Strikingly, we found that both AT-hooks were unable to stably bind DNA at the same time (Fig. 4D, Fig. S8A). Rather, individual AT-hooks switched between binding to DNA and binding to NCR1 (Fig. 4D, Fig. S8A). In contrast, we found that an N-terminal D1 peptide containing 3 AT-hooks uninterrupted by negative charges (hereafter DNA-binding module 1 or DBM1) permitted simultaneous binding of all AT-hooks to DNA (Fig. 4E, Fig. S8B), which led to a significant increase in DNA-binding strength (Fig. 4F, Fig. S8C), likely due to cooperative binding of the AT-hooks. Based on these data, we designated 4 DNA-binding modules (DBMs) on D1, each consisting of 2-4 AT-hooks, which are interspersed by 4 NCRs. Interestingly, further MD simulations revealed the potential for direct interactions between DBMs and NCRs, largely driven by positive charges on the AT-hooks and negatively charged residues on NCRs (Fig. 4G, Fig. S8D, E). Taken together, our molecular dynamics simulations predict that D1 consists of 4 DBMs that can directly bind DNA with an interaction valency of 4. At the same time, the DBMs can also associate with NCRs, potentially mediating D1 self-association.

**Figure 4.**
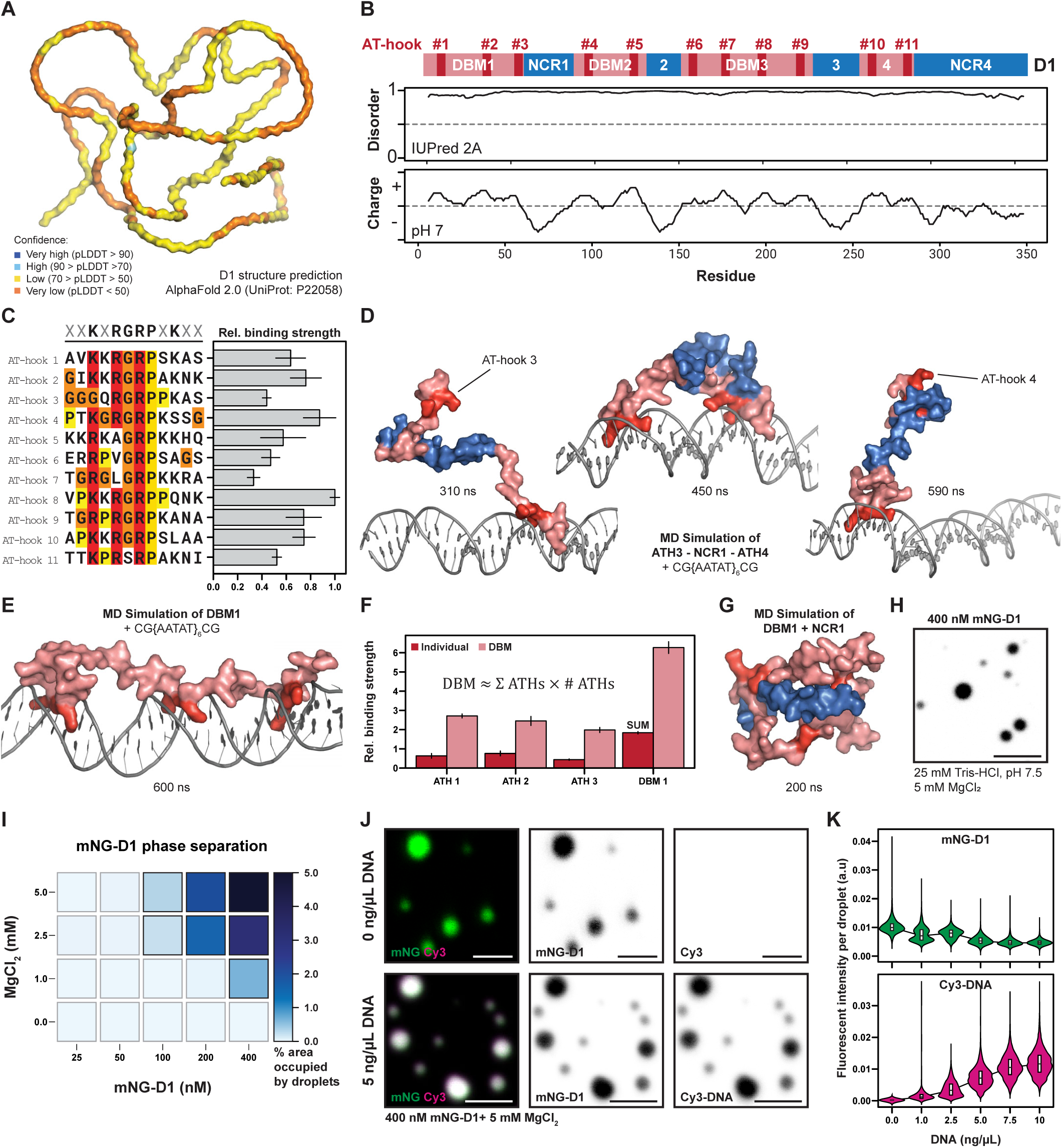
D1 phase separates through a complex coacervation-like process. (A) AlphaFold 2.0 prediction of D1 structure. (B) Overview of D1 sequence features, showing the location of its 11 AT-hooks, charge-based modules and predicted disorder. (C) Sequence alignment of the 11 D1 AT-hooks and predicted DNA binding strength from steered MD simulations. (D) MD simulation snapshots of the ATH3-NCR1-ATH4 peptide interacting with DNA. (E) Bound conformation of DBM1 with DNA. (F) Predicted relative binding strength of DBM1 vs. AT-hooks #1-3. Red bars represent results from simulations of individual AT-hooks, pink bars show data from DBM1-DNA simulation. (G) MD simulation snapshots of DBM1 and NCR1 (unconnected peptides). (H) Fluorescent microscopy image of mNG-D1 in minimal buffer with MgCl2. (I) Phase diagram of mNG-D1 at different MgCl2 concentrations. Phase separation was quantified as fraction of area covered by droplets. Black outlines indicate presence of phase separation. (J) Fluorescent microscopy images of mNG-D1 and Cy3-labelled AATAT plasmid at different DNA concentrations. Scale bars: 5 µm. (K) Fluorescent intensity of mNG-D1 and Cy3-AATAT plasmid within the droplets at different DNA concentrations. Bar plots display bootstrapped mean and standard deviation (see materials and methods).

We next set out to test the DNA-binding and predicted self-association of D1 using purified protein. First, we used an electrophoretic mobility shift assay (EMSA) to assess the ability of D1 to bind a series of 60bp oligonucleotides, each consisting of different 5bp repeats, *in vitro* (Fig. S9A). We found that D1 bound a 12xAATAT oligonucleotide with ∼3-fold higher affinity in comparison to a control 60bp non-repetitive oligonucleotide (Fig. S9B, C). D1 was also able to bind other AT-rich repeat sequences such as 12xAATAG and 12xAATAC with relatively high affinity (Fig. S9C). We observed that repeat tract length influenced D1 binding to DNA, with a steady increase in affinity observed for 60bp oligonucleotides containing 4xAATAT, 8xAATAT and 12xAATAT (Fig. S9C, D). Overall, these data are an extension of previous reports on the binding affinity of D1 to AT-rich satellite DNA tracts in nuclear extracts^68,69^ and are consistent with the fact that D1 clusters Mb-length tracts of AATAT into chromocenters *in vivo*^41^.

Based on the predicted charge interactions between DBMs and NCRs, we next assessed whether D1 was able to self-associate *in vitro*. We observed that D1 formed two distinct phases in a minimal buffer system containing MgCl_2_ at sub-physiological concentrations as low as 100nM (Fig. 4H, I). Given the charged nature of the DBMs and NCBs, we posit that D1 self-association occurs through complex coacervation, an electrostatic interaction-driven phase separation process. Since chromocenters contain both D1 and DNA, we next used a Cy3-labelled 5kb plasmid containing a ∼ 1kb AATAT tract to test the effect of DNA on D1 phase separation. At low concentrations, we found that the plasmid DNA was able to effectively partition into D1 condensates (Fig. 4J, K). However, increasing amounts of DNA led to a reduction in the amount of D1 contained within the condensates, likely due to the competition between DNA and NCRs for DBM binding (Fig. 4K). Consistently, the addition of excess DNA (100ng/µl and greater) was sufficient to abolish D1 phase separation. These data further imply that electrostatic interactions are important for D1 phase separation *in vitro* and may underlie the clustering of D1-bound DNA into chromocenters *in vivo*.

### A modular charge pattern is required for D1-dependent chromocenter formation in vivo

We have previously demonstrated that ectopically expressed D1 in cultured mouse cells localizes to chromocenters and enhances their coalescence in a concentration-dependent manner^41^. To better understand the dynamics of this process, we performed live imaging of cultured mouse NIH-3T3 fibroblasts expressing mNeonGreen(mNG)-tagged D1^WT^. Excitingly, we observed chromocenter fusions that occurred in the order of minutes (Fig. 5A, arrowheads indicate fusing chromocenters) and occasional fissions that occurred on a slightly longer time scale (Fig. S11A), reinforcing the notion that chromocenters are satellite DNA-containing biomolecular condensates *in vivo*^38–40^. Over longer time periods, we observed a consistent decrease in the number of chromocenters per nucleus, in a manner that primarily relied on D1 expression levels (Fig. 5B-D). However, our live imaging data also revealed a potential role for cell movement in regulating chromocenter fusion (Fig. S11B, C), likely due to motility-induced changes in nuclear organization that promote encounters between individual chromocenters.

**Figure 5.**
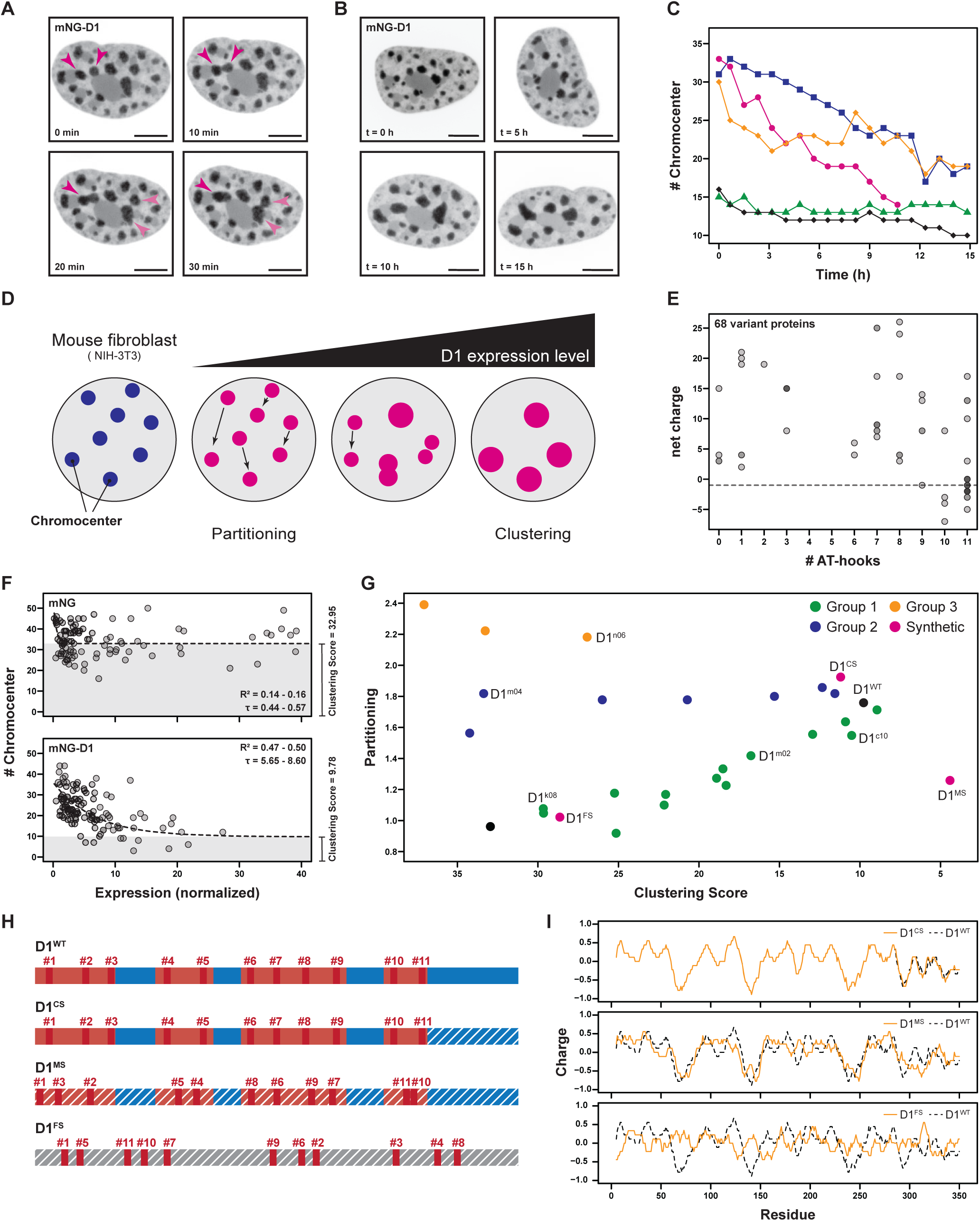
Multivalency and charge modules on D1 are required for satellite DNA clustering into chromocenters. (A) mNG-D1 positive chromocenters in NIH-3T3 nuclei undergoing fusion. (B) Chromocenters in a NIH-3T3 nucleus expressing mNG-D1 over the course of 15 hours. (C) Quantification of number of chromocenters over 15 hours in 5 NIH-3T3 cells expressing mNG-D1. Each color represents a different cell (D) Schematic showcasing the effect of expression of D1 in mouse cells; D1 (red) partitions into pre-existing mouse chromocenters (blue) and clusters them into fewer, larger chromocenters as D1 expression levels increase. (E) Number of AT-hooks and net charge of proteins within the D1 variant library. (F) Number of chromocenters in relation to normalized expression in fixed NIH-3T3 cells 48 hours post transfection of mNG and mNG-D1^WT^. The dashed line indicates the median bootstrapped fit. (G) The partitioning and chromocenter clustering of selected D1 variants in NIH-3T3 cells 48h post transfection. (H) Schematic of D1^CS^ (C-terminal Scramble), D1^MS^ (Module Scramble), and D1^FS^ (Full Scramble). Dashed lines indicate scrambled region on D1. (I) Charge profiles of D1 scramble variants. Scale bars: 5 µm.

Thus far, our *in vitro* results suggest that D1 phase separation may be critical for clustering D1-bound satellite DNA repeats into chromocenters. We set out to test this idea in a cellular context by generating a panel of D1 variants using a combinatorial CRISPR/Cas9-based approach in the *Drosophila* germline (Fig. S10A). Notably, this approach yielded a panel of 68 fly strains carrying unique mutations at the endogenous D1 locus, with these D1 variants exhibiting a wide range of net charge and AT-hook numbers (Fig. 5E, Fig. S10B). To rapidly and quantitatively determine the functional properties of the D1 variants, we leveraged a previously developed assay in mouse cultured cells ^41,70^. Briefly, we expressed mNG, mNG-D1^WT^, as well as 24 of the most distinct D1 variants (Fig. S10B), in cultured mouse NIH-3T3 fibroblasts and analyzed the number of chromocenters in fixed cells 48h post transfection. We used 3D segmentation to quantify the volumes of whole nuclei as well as individual chromocenters and employed fluorescent beads to normalize expression levels across different cells, replicates and D1 variants (see materials and methods). NIH-3T3 fibroblasts contain ∼75 chromosomes^71^, which are normally clustered into 30-35 chromocenters. We found that expression of mNG alone did not change the number of chromocenters (Fig. 5F). On the other hand, mNG-D1^WT^ expression caused a concentration-dependent decrease in the number of chromocenters, which we fitted using a mono-exponential decay function (Fig. 5F). We used bootstrapped curve fitting to extract a clustering score (the horizontal asymptote) which represents the expected minimum number of chromocenters upon infinite protein expression. We also estimated the partitioning coefficient, which quantifies the extent to which the expressed protein localizes into chromocenters. Expression of mNG was associated with a partitioning co-efficient of 0.96 and a median clustering score of 32.95 (Fig. 5F, G). In marked contrast, D1^WT^ exhibited a partitioning co-efficient of 1.76 and a median clustering score of 9.78 (Fig. 5F, G), consistent with its known ability to localize to mouse chromocenters and enhance their coalescence in an expression-dependent manner. We performed the same analysis for each of the 24 D1 variants (Fig. S10B, Fig. S12), allowing us to determine the effect of charge, modularity, and multivalency on D1’s chromocenter clustering function (Fig. 5G). Based on these data, we separated the D1 variants into three functional groups which differed in their ability to partition and coalesce chromocenters. The D1 variants in group 1 exhibited a linear relationship between partitioning and coalescence. A closer examination of these variants revealed that their coalescence and partitioning was dependent on both the number of AT-hooks and net charge (Fig. S13A-D). In contrast to the group 1 D1 variants, we found that group 2 D1 variants exhibited differential coalescence ability even though their partitioning coefficients were relatively similar. The unaffected partitioning into chromocenters is most likely explained by the fact that the group 2 variants retain at least 7 AT-hooks (Fig. S13A-D). However, we observed a strong correlation between positive net charge in group 2 variants and defective coalescence (Fig. S13B). Group 3 D1 variants, which retain between 9-11 AT-hooks, partitioned even more strongly into chromocenters than group 2 variants but exhibited poor clustering scores, again likely due to their significant positive net charge (Fig. S13A-D). Our data suggest that the number of AT-hooks on D1 is the main determinant of chromocenter partitioning while a near neutral net charge appears to be critical for chromocenter coalescence in mouse nuclei.

While our data point towards a model where electrostatic interactions between D1 DBMs and NCRs largely drive chromocenter coalescence, we were still not able to exclude potential contributions by other factors such as unidentified protein-protein interaction motifs. Therefore, we designed 3 synthetic D1 constructs to further dissect the key determinants of D1-dependent chromocenter coalescence (Fig. 5H, I). First, we designed a D1 C-terminal scrambled (D1^CS^) variant, which scrambles the NCR4 sequence (40% sequence identity) but maintains the charge pattern (Fig. 5H, I). We also designed a module scrambled (D1^MS^) variant, which randomizes residues within each module (i.e. DBMs and NCRs) but retains the overall charge pattern. Finally, we designed a full scrambled (D1^FS^) variant, which retains 11 AT-hooks but lacks the modular organization and charge pattern of D1. In all the synthetic variants, the 5 core residues of each AT-hook (XRGRP) were kept intact during sequence scrambling to maintain DNA-binding. When expressed in mouse cells, we found that the D1^CS^ and D1^MS^ variants exhibited a D1^WT^-like ability to induce Chromocenter coalescence (Fig. 5G). In marked contrast, the D1^FS^ variant did not preferentially partition into chromocenters and did not enhance coalescence (Fig. 5G). Taken together, these data emphasize that the modular charge pattern (i.e. DBMs and NCRs) is a critical factor for D1-dependent chromocenter coalescence *in vivo*. We propose that satellite DNA condensation into chromocenters occurs through (a) the ability of D1 to selectively bind satellite DNA repeats across multiple chromosomes and (b) the phase separation of DNA-bound D1 molecules.

### Weakened chromocenter compaction and coalescence impairs nuclear mechanostability

The generation of fly strains carrying the individual D1 variants characterized above allowed us to test the effect of chromocenter compaction and coalescence on nuclear mechanostability *in vivo*. We selected 5 D1 variant strains (Fig. 6A), which exhibited a broad range of clustering scores in mouse fibroblasts (Fig. 5G, Fig. S11B) and performed an in-depth analysis of their effect on chromocenter formation and nuclear mechanostability in *Drosophila* spermatogonia. We observed that spermatogonia containing three of the selected variants (D1^k08^, D1^n06^ and D1^c10^) exhibited de-compacted chromocenters (increased AATAT volume fraction, Fig. 6B), which was associated with nuclear deformation (Fig. 6C). Interestingly, D1^c10^-containing mouse chromocenters were irregularly shaped and occasionally thread-like (Fig. S14A, B). Therefore, despite exhibiting D1^WT^-like clustering in mouse cells, D1^c10^-containing spermatogonia cannot maintain satellite DNA condensation in the mechanically challenged *Drosophila* testis. Together, these three D1 variants highlight the importance of chromocenter compaction for nuclear mechanostability.

**Figure 6.**
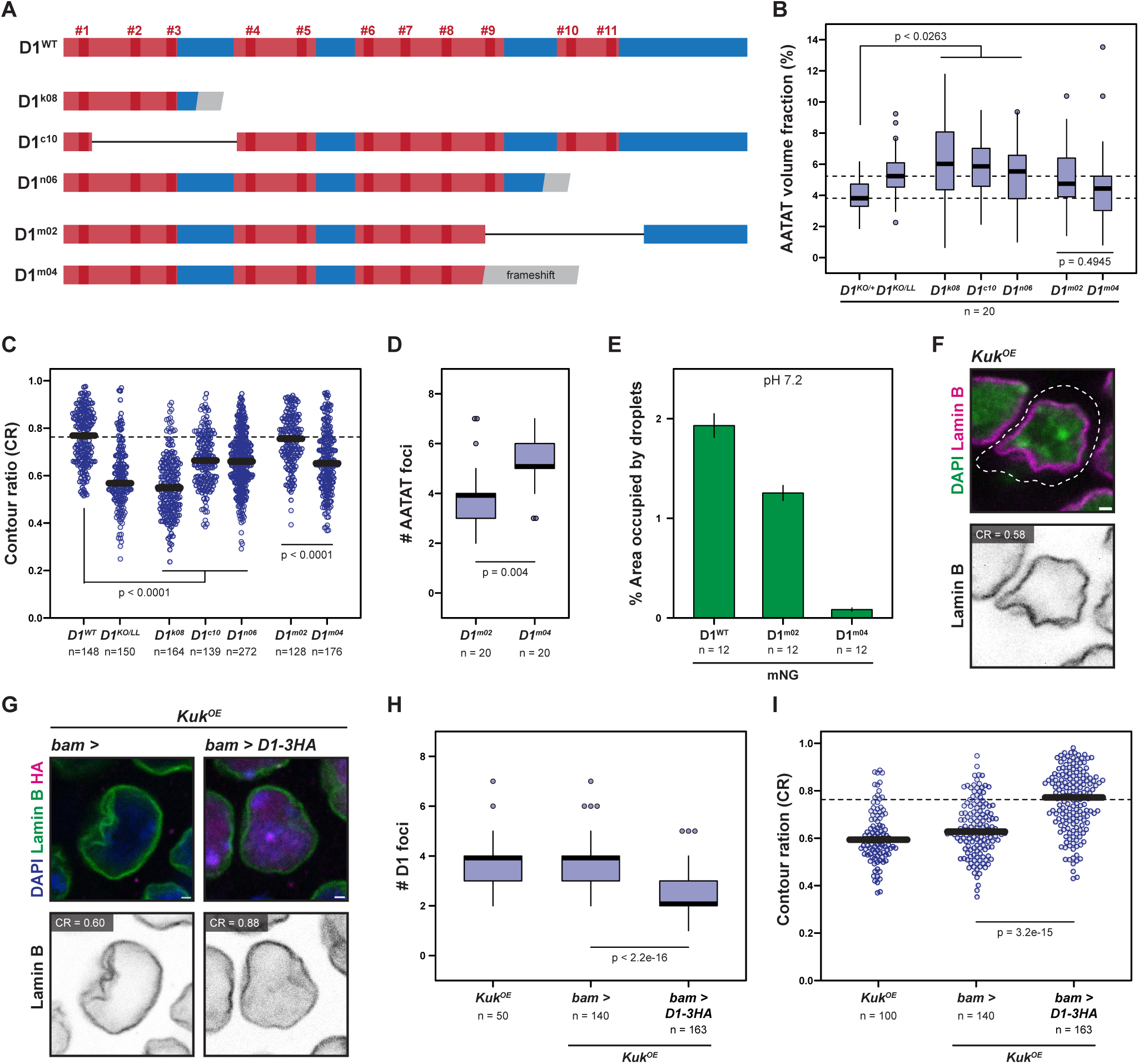
Chromocenter formation is a tuneable nuclear mechanostability mechanism in *Drosophila* spermatogonia. (A) Schematic of selected D1 variants analyzed in *Drosophila* spermatogonia. (B) AATAT volume fraction in adult spermatogonia from the indicated genotypes. *D1^KO/+^* and *D1^KO/LL^* datapoints replotted from Fig. S1E. Dashed lines represent median values from *D1^KO/+^* and *D1^KO/LL^* spermatogonia. (C) CR for adult spermatogonia expressing the indicated D1 variant proteins. (D) Number of AATAT foci per nucleus in adult D1^m02^ and D1^m04^ spermatogonia. (E) Phase separation of D1^WT^, D1^m02^ and D1^m04^ at pH7.2. Error bars show the standard error of the mean. (F) Kuk^OE^ spermatogonia stained for Lamin B (magenta) and DAPI (green). (G) Spermatogonia from the indicated genotypes stained for Lamin B (green), HA (magenta) and DAPI (blue). (H, I) Number of D1 foci (H) and CR (I) in adult spermatogonia from the indicated genotypes. Indicated p-values are from Mann-Whitney U-tests. For the CR plots, the thick black line indicates the median while the dashed line represents the median CR (0.76) from adult *D1* heterozygous spermatogonia. All scale bars: 1 µm.

We next focused our attention on two D1 variants containing 8 AT-hooks, D1^m02^ and D1^m04^ (Fig. 6A). These two variants are primarily differentiated by their net charge; while NCR4 is present in D1^m02^ and contributes to an overall neutral charge, D1^m04^ contains a positively charged frameshift at its C-terminus. Importantly, we have also demonstrated that a positive net charge impairs chromocenter coalescence in mouse nuclei. While neither variant affected satellite DNA compaction (AATAT volume fraction, Fig. 6B), we noted that the number of AATAT-containing chromocenters in D1^m04^-containing spermatogonia was significantly higher than in D1^m02^-containing spermatogonia (Fig. 6D). To better understand the mechanistic basis for defective chromocenter coalescence by D1^m04^, we turned to *in vitro* analysis of purified proteins. Both mNG-D1^m02^ and mNG-D1^m04^ exhibited DNA-binding that was at the same level or improved in comparison to mNG-D1^WT^ (Fig. S9E). However, D1^m02^ phase separation was significantly stronger than D1^m04^ at a near neutral pH (Fig. 6E) and likely underlies the more effective chromocenter coalescence *in vivo*. Importantly, D1^m04^-containing spermatogonia also exhibited increased nuclear deformations in comparison to spermatogonia expressing the more effective D1^m02^ variant. Analysis of additional variants (Fig. S14C) revealed a positive correlation (R^2^ – 0.60) between D1 variant clustering scores in mouse cells and nuclear mechanostability in *Drosophila* spermatogonia (Fig. S14D). Overall, we conclude that satellite DNA condensation into chromocenters is a key determinant of nuclear mechanostability in a tissue context.

### Enhanced chromocenter coalescence improves nuclear mechanostability

To complement our previous experiments, we attempted to determine whether enhanced chromocenter coalescence could improve nuclear mechanostability. To do so, we turned to the *Drosophila* inner nuclear membrane protein Kugelkern (Kuk), which has been linked to the deformability of the nucleus^72–77^. Loss of Kuk results in more rigid^75^ and less deformable nuclei^72–77^ while Kuk overexpression induces nuclear deformations across yeast, *Drosophila*, *Xenopus* and mouse^72,73^. In spermatogonia, we found that Kuk overexpression (Kuk^OE^) in an otherwise WT background led to increased nuclear deformations (median CR = 0.59), without affecting the number of chromocenters (Fig. 6F, H, I). We next assessed whether enhancing chromocenter coalescence would modify Kuk^OE^ - dependent nuclear deformations. We found that D1 overexpression (D1^OE^) led to a reduced number of D1 foci in spermatogonia i.e. enhanced chromocenter coalescence (Fig. 6H). Excitingly, nuclear deformations were also reduced to WT levels when both D1 and Kuk were overexpressed in spermatogonia, in comparison to the Kuk overexpression alone (Fig. 6G, I). Our data show that enhancing satellite DNA condensation into chromocenters can improve the ability of nuclei to withstand mechanical stress. Taken together, we propose that nuclear mechanostability can be tuned by modulating the degree of satellite DNA condensation into chromocenters.

## DISCUSSION

Satellite DNA repeats are often present as 10^5^-10^7^ bp length tracts at the centromeric and pericentromeric heterochromatin of eukaryotic chromosomes^33^. Despite their abundance, these non-coding repeats have been historically dismissed as ‘junk DNA’, in part due to lack of clear discernible function(s)^78,79^. Therefore, whether satellite DNA propagate selfishly, are maintained passively or actively contribute to cellular function and evolutionary fitness remains actively debated. In this study, we use a combination of experiments and *in silico* modeling to demonstrate that the condensation of satellite DNA repeats into chromocenters gives rise to nuclear mechanostability. In the *Drosophila* testis, a tissue subject to persistent mechanical stress, we found that chromocenter disruption is strongly associated with nuclear deformations in spermatogonia while enhancing chromocenter coalescence in the same cells improves nuclear mechanostability. We propose that chromocenters represent physical connections between different chromosomes that allow the genome to function in unison against mechanical stress (Fig. 7). The conservation of chromocenters across insects, plants and mammals suggests that these nuclear condensates contribute to mechanostability and genome integrity across a wide range of mechanically challenged cells and tissues. As the lynchpins of chromocenter formation, satellite DNA repeats justify their existence in eukaryotic genomes.

**Figure 7.**
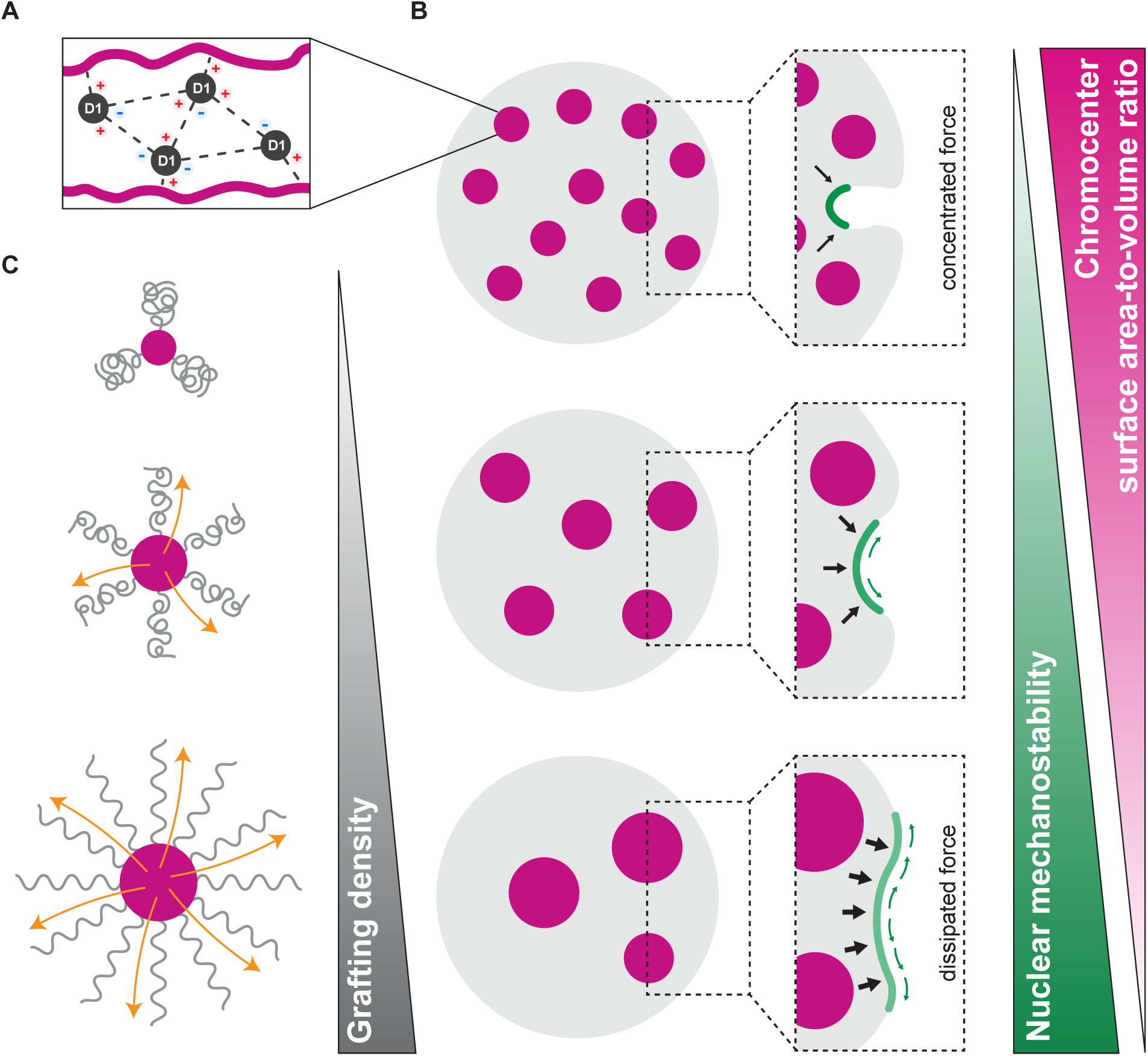
Condensation of satellite DNA from multiple chromosomes into chromocenters tunes nuclear mechanostability. (A) Charge-based DBM-NCR interactions between D1 molecules underlie the condensation of D1-bound satellite DNA repeats into chromocenters. (B) Enhanced satellite DNA condensation (i.e. fewer and larger chromocenters) scales with nuclear mechanostability, likely through dissipation of the applied force over a larger nuclear surface (green region) due to increased internal pressure (black arrows). (C) Chromocenter coalescence reduces the surface area-to-volume (SA:V) ratio and the corresponding increased grafting density of the surrounding chromatin may increase outward pressure (orange arrows).

Interestingly, the polymer modeling and theory suggest that the degree of chromocenter coalescence increases internal nuclear pressure such that external compressive forces are dissipated over a larger nuclear envelope surface (Fig. 7B). We suggest that the chromocenter-dependent nuclear pressure partly originates from physically linked chromosome ensembles being less prone to displacement by mechanical stress, especially in comparison to individual chromosomes. A similar reduction in chromatin displacement has been proposed as the mechanism underlying LINC complex-based nuclear mechanostability^10^. In addition, increasing the degree of chromocenter coalescence reduces the surface area-to-volume (SA:V) ratio of these condensates (Fig. 7B) and we speculate that this phenomenon may also play a significant role in nuclear mechanics. For example, interphase chromosomes have been modelled as polymer-grafted colloidal particles, where ‘rigid cores’ consisting of chromocenters are grafted by polymers consisting of chromatin loops from the rest of the chromosome^80^. We propose that in nuclei containing many chromocenters, the relatively higher chromocenter SA:V ratio is associated with a lower grafting density of non-satellite DNA chromatin, which likely adopts an energetically favorable compact conformation (Fig. 7C). As chromocenters coalesce, we posit that the reduced SA:V ratio leads to an increased grafting density of the surrounding chromatin, changing its conformation to a more extended form and thus increasing outward pressure (orange arrows, Fig. 7C). In this model, the chromocenter surface tension, potentially regulated by viscoelasticity, may be critically important to allow the surrounding chromatin in ‘pushing back’ against mechanical compression.

Given the range of constitutive heterochromatin organization in eukaryotic nuclei, we propose that chromocenters likely function in concert with other heterochromatin features such as histone modifications, peripheral tethering and cross-linking, with the relative contribution of each of these features dependent on the mode of heterochromatin organization in the specific cell type or organism. We also note that not all satellite DNA-containing species (e.g. humans and certain plants) exhibit prominent DAPI-dense chromocenters^33,81^. However, recent reports demonstrate that systematic detachment of constitutive heterochromatin from the nuclear periphery in human cells leads to the formation of DAPI-dense chromocenter-like foci^31,32^. Similar heterochromatin foci, known as senescence-associated heterochromatin foci (SAHF), form due to the loss of peripheral heterochromatin attachments during senescence and require the D1 functional orthologue, HMGA1^82,83^. Interestingly, chromocenter-like foci are absent in diseases such as progeria and other laminopathies, which also exhibit reduced peripheral heterochromatin^84,85^. We speculate that the lack of chromocenters may contribute to the deleterious nuclear deformations and DNA damage observed in these pathologies^84–87^. Therefore, physically connecting chromosomes through satellite DNA repeats may be an underappreciated mechanism ensuring nuclear mechanostability and genome integrity, even in organisms that do not typically exhibit chromocenters.

## ACKNOWLEDGMENTS

We thank members of the Jagannathan lab, Hugo Stocker and Yves Barral for discussion and comments on the manuscript. We are grateful for the reagents and resources provided by Yukiko Yamashita, Bernhard Hampoelz, Yikang Rong, Tzumin Lee, the Bloomington Drosophila Stock Center, the Vienna Drosophila Resource Center, the Kyoto Drosophila Stock Center and the Developmental Studies Hybridoma Bank. We acknowledge microscopy support from the Scientific Center for optical and Electron Microscopy (ScopeM), NGS support from the Functional Genomics Center Zurich (FGCZ) and bioinformatic infrastructure from Shinichi Sunagawa. We acknowledge the following funding sources, Swiss National Science Foundation (310030_189131 and 320030_228043) – MJ Swiss National Science Foundation (310030_207588 and 310030_205199) – PC European Research Council (101018257) – PC European Research Council (101002094) – PA Dutch Research Council (OCENW.KLEIN.200) – JV

## AUTHOR CONTRIBUTIONS

Conceptualization, F.B.B., B.F., and M.J.; methodology, F.B.B., B.F., I.C., L.d.M., T.v.H., A.S.; Investigation, F.B.B., B.F., I.C., L.d.M., T.v.H., A.S., F.R., S.S.L., C.M., A.S.; writing—original draft, F.B.B., B.F., and M.J.; writing—review & editing, F.B.B., B.F., I.C., L.d.M., T.v.H., C.M., P.A., A.S., T.C.T.M., P.C., J.V., and M.J.; funding acquisition, P.A., P.C., J.V., and M.J..; supervision, T.C.T.M., P.C., J.V., and M.J.

## MATERIALS AND METHODS

### Fly husbandry and strains

All flies were raised on standard Bloomington medium at 25 °C unless stated otherwise. Crosses were performed at 25 °C and 70 % relative humidity. *D. melanogaster y w* was used as a wild-type strain. *koi*^80^ (BDSC25105), *UAS-bam^RNAi^* (BDSC58178), *UASt-NLS-GFP* (BDSC4776), *UAS-Lamin B-GFP* (BDSC7377), *UAS-Lamin B^RNAi^* (BDSC57501), *UAS-HP1^RNAi^* (BDSC36792), *UAS-Kuk^RNAi^* (BDSC60449) and *CenpC-tdTomato* (BDSC91717) were obtained from the Bloomington Drosophila Stock center. D1LL03310 (DGRC140754) was obtained from the Kyoto Stock Center. *EGFP-aub* (VDRC313243) was obtained from the Vienna Drosophila Research Center. *nos-Gal4* (Chr. 2) and *bam-Gal4* were gifts from Yukiko Yamashita. *EGFP-HOAP* was a gift from Yikang Rong.

*Kuk-EGFP* was a gift from Bernhard Hampoelz and Martin Beck. *UAS-D1-3xHA* was a gift from Daniel Barbash. *bam-Cas9* was a gift from Tzumin Lee. *UASt-GFP-D1* has been previously described^41^. Clones bearing D1-positive and D1-negative cysts were generated by crossing *hsFlp; FRT82B, nls-RFP* to *FRT82B, D1^KO^/ TM6B* and collecting progeny for 24 to 48 h. Mitotic clones were induced by subjecting the late embryos/early larvae to a 1h heat shock at 37 °C. The progeny were allowed to develop to the 3° instar larval stage or adulthood at 25 °C prior to dissection.

### Generation of transgenic strains and mutant alleles

The D1knockout (*D1^KO^*) allele was generated by replacing the D1 ORF with a DsRed cassette using CRISPR-mediated homology-directed repair. Briefly, 700-1000bp from the 3′ UTR and 5’ UTR were cloned into a vector (pBSK-attB-DsRed-attB) flanking a 3xP3-driven DsRed cassette. This plasmid was coinjected along with two gRNA-expressing plasmids (pU6-Bbs1-chiRNA containing gRNA1: GAAGTTGCGGTAAAGAAGCG and gRNA2: ATCCGAAACATAACCGTCGT) in embryos from the nos-Cas9 strain (BDSC78781) by BestGene, Inc.

Transformants were selected based on DsRed expression, proper integration into the *D1* locus was verified by PCR, and resulting flies were outcrossed into a wild type background for 6 generations before use. For the *UASt-GFP-HMGA1* transgene, the HMGA1 cDNA was synthesized (Twist) and subcloned into pUASt-GFP-attb^88^. Transgenic flies were generated by PhiC31 integrase-mediated integration into the *attP40* site (BestGene). For the *D1* variant alleles, we used *bam-Cas9* and a variety of gRNA strains (Fig. S11A, Supplementary Table 1) to introduce mutations at the endogenous locus. gRNAs were picked using the CRISPR Optimal Target Finder (http://targetfinder.flycrispr.neuro.brown.edu/) and a CRISPR Efficiency Predictor tool (https://www.flyrnai.org/evaluateCrispr/). Candidate gRNAs with a score below 7.4 were discarded. A final set of 8 gRNAs was chosen based on low off-target effects. An overview of the gRNA sequences used in this study is provided in Supplementary Table 1. For further details on the variants and gRNA sequences refer to the associated repository. Subsequently, bam-Cas9 flies were crossed to the following gRNA strains: (i) a single gRNA strain targeting the D1 sequence (BDSC:84068; D1 variants starting with c, e, and h), (ii) strains with two gRNAs using the pCFD4 plasmid (D1 variants starting with j-o) and (iii) randomly selected gRNA pairs based on the Cas9-mediated Arrayed Mutagenesis of Individual Offspring (CAMIO) approach (D1 variants starting with i). Overall, 68 lines carrying unique edits in the D1 sequence were established and characterized via PCR amplification of the genomic D1 sequence and Sanger sequencing (Microsynth AG, CH).

### Immunofluorescence staining, microscopy and image analysis in Drosophila tissues

For immunofluorescence microscopy of *Drosophila* tissues, larval or adult testes were dissected in 1x PBS and transferred to 4% paraformaldehyde in PBS for a 20 min fixation at room temperature on a nutator. The samples were then washed three times for 20 min each in 1x PBS containing 0.1% Triton-X (PBS-T) before blocking in 3% BSA in 1x PBS-T for at least 30 minutes. Primary antibodies were added and incubated with the samples at 4°C overnight. Samples were again washed three times with PBS-T, blocked and incubated with secondary antibodies in 3 % BSA in PBS-T either overnight at 4°C or 3 to 5 h at room temperature. Finally, the samples were washed as above and mounted on glass slides in VECTASHIELD® Antifade Mounting Medium with DAPI (Vector Laboratories, USA). The following primary antibodies were used in this study: mouse anti-Lamin Dm0 (1:200; ADL84.12, DSHB), rat anti-Vasa (1:100; AB_760351, DSHB), rabbit anti-γH2Av (1:500; pS137, Rockland), rabbit anti-HA-tag (1:400; ab9110, Abcam), mouse anti-HP1 (1:50; C1A9-s, DSHB). Guinea pig anti-Otefin (1:200) was a gift from Georg Krohne. Rabbit anti-Vasa (1:200) was a gift from Yukiko Yamashita. Rabbit anti-Prod (1:200) was a gift from Tibor Torok. Guinea pig anti-D1 (1:500) has been previously described^41^. The rabbit anti-γH2Av antibody was pre-adsorbed in 3% BSA in PBS-T over night on 5 fixated ovaries before using it for testes staining. Fluorescent microscopy images (z-stacks) were acquired using a Leica TCS SP8 confocal microscope with 63x oil-immersion objectives (NA = 1.4). Images were processed using ImageJ/FIJI. Briefly, for each testis, a plane with the maximum number of visible spermatogonial nuclei was used to select the cohort of cells for analysis. To determine the micronucleation rate, all cells in the cohort were assessed for the presence of micronuclei in 3D. For the nuclear contour ratio (CR), 10 to 15 spermatogonia from the 8-16 cell stage were chosen randomly from the cohort. The CR was obtained by empirically determining the z-plane of maximal nuclear envelope deformation for each cell individually followed by a measurement of the nuclear perimeter and area in FIJI. The CR was then calculated as 4π x area/perimeter^2^ to represent the amount of deformation per nucleus. Where applicable, RNAi knockdown efficiencies were quantified by extracting the mean gray value of the respective fluorophores or antibody staining per nucleus. The values were normalized to the mean intensity of a subset of cells in the same tissue that were unaffected by the knockdown (e. g. somatic cells). For live imaging, testes of larval and adult flies expressing Aub-GFP were dissected into 1x Schneider’s Drosophila medium (Gibco) pre-warmed to 25°C. They were placed in a glass bottom dish, weighed down with a cropped dialysis membrane (Spectra/Por, Spectrumlabs) and immediately imaged on a Leica TCS SP8 confocal microscope (400 Hz, 10s, 50 frames). Movement was quantified in ImageJ/FIJI by manually tracking bright Aub-GFP foci per frame. 5 tracks were recorded per testis.

### DNA FISH and volume fraction measurements

After immunofluorescence staining, samples were post-fixed with 4% formaldehyde for 10 min and washed in PBS-T for 30 min. Fixed samples were incubated with 2 mg/ml RNase A solution at 37°C for 10 min, then washed with PBS-T. Samples were denatured at 95 °C for 5 min and rinsed with 2x SSC-T (2xSSC containing 0.1% Tween-20). Hybridization buffer (60% formamide, 10% dextran sulfate, 2x SSC, 1 µM probe) was added to the samples and incubated overnight at room temperature. The following fluorescent probes were used 488-(AATAT)_6_, Cy3-(AATAT)_6_, Cy3-(AATAC)_6_, Cy5-(AGG ATT TAG GGA AAT TAA TTT TTG GAT CAA TTT TCG CAT TTT TTG TAA G), Cy5-(ACC GAG TAC GGG)_6_, Cy3-(AAGAT)_6_, Cy5-(AATAGAC)_5_, Cy5-(TAGA)_7_. Images were processed using ImageJ/FIJI. The chromocenter volume fraction was obtained by manually segmenting nuclei and chromocenters in 3D.

### Fabrication of tweezers for manual force application

Tweezers with a defined cavity of 140 µm were manufactured using SMD-tweezers with an initial cavity diameter of 0.6 mm (SM116.SA.1, ideal-tek).

Briefly, 2-component-epoxy glue (UHU PLUS 2-K-Epoxidkleber schnellfest) was mixed in a 1:1 ratio, filled into the cavity and incubated at room temperature for roughly 5 min until the epoxy hardened a bit but remained moldable. The tweezers were then dipped in oil, and a piece of plastic wrap was attached at each side of the tips to prevent the wire from sticking to the epoxy. A wire with a diameter of 140 µm was positioned between the tips, the tweezers were closed firmly and secured by tightly wrapping tape around the base to keep it closed. The product was then kept at 60°C on a heat block for 30 min covered in aluminum foil and left overnight at room temperature to fully harden. After removal of the tape, wrap, wire and oil, the tips were cleaned with PBS. To re-fabricate a tweezer, the hardened epoxy can be carefully removed with a razor blade. Larval testes were dissected in 1x PBS and compressed with the tweezers 20 times over the course of 15-20 seconds. They were then fixated in 4 % paraformaldehyde in PBS for 20 min and processed as described above.

### Coarse-grained nuclear model

A simplified nuclear model^58^ was adapted and simulated using the HOOMD-blue simulation engine v4.7.0. Briefly, chromatin monomers were placed on a grid at bond-length distance (*l*_*chrom*_) using the grow_cubic function from polykit (https://github.com/open2c/polykit) to achieve a densely packed initial chromatin conformation. 20 chromosomes consisting of 1000 particles each were placed in a simulation box in this manner. The first 100 particles of each chromosome were designated as ‘satellite DNA’ to allow force field parameters to be varied for this subset of chromatin particles. A final shell radius (*R*_*final*_) was determined based on the number of chromosome monomers, aiming for a final chromatin volume fraction of 25%. Shell particles were placed on a sphere with a radius of *R*_*initial*_ = 4/3 ⋅ *R*_*final*_ to ensure encapsulation of the initial chromatin conformation, and connections were defined using the ConvexHull function within the scipy package (version 1.13.1). North pole (NP) and south pole (SP) particles were designated from opposite sides of a randomly assigned squeezing-axis. Chromatin positions were frozen during initial energy minimization to achieve relaxation of the initial shell conformation to *R*_*final*_. SP particle coordinates were frozen from this point throughout the simulation. Attractive forces between ‘satellite DNA’ particles were set to either 0.0ε, 0.05ε, 0.1ε, 0.2ε, or 0.4ε (ε = *k*_*B*_*T*), and an initial 1e6 simulation steps were run using a short integration step size (5e-6), followed by the 1e8 step production run at the standard step size of 5e-4. We simulated the mechanical challenge following the equilibration runs by placing a wall potential tangential to the south pole and pushing the NP particle towards the SP 20 times, allowing for relaxation of the system between ‘squeezes’. Specifically, a constant force was applied to the NP towards the SP using the normalized vector between the two points (|*v*→_*sq*_| = 1), multiplied with the force constant *F*. *F* was increased linearly from 0.01 to 100 every 5e5 simulation steps in 10 iterations, where *v*→_*sq*_ was updated with each increment of *F*. Each compression was followed by a relaxation run of 2e6 steps. Production runs were carried out on the ETH Euler cluster. Analyses were carried out using custom Python scripts accessible via the associated repository.

Chromocenters were defined during the simulations as non-overlapping sets of chromosomes making non-bonded contacts via their ‘satellite DNA’ particles-contacts were defined via a distance cutoff set to 2*l*_*chrom*_. The bond-tension of shell particles was calculated via bond-length deviations from the resting bond length (Δ*l*_*shell*_). Bond-length deviations were converted into energies via the force field equation *E*_*shell*_ = ^1^ *k*_*shell*_ ⋅ Δ*l*_*shell*_, where the harmonic constant *k*_*shell*_ = 15ε. Shell particles were converted into a graph using networkx (version 3.2.1), and median energies were grouped by the bond-distance from the NP. A baseline tension cutoff was defined at 3 standard errors from the global tension median, and tension persistence was defined as the point where the window-averaged tension profile recovers to this baseline (as illustrated in Fig. 3F). Visualizations were created using VMD^89^.

### Atomistic MD simulation

A reference AT-hook structure (PDB ID: 2ezf, ^90^) was placed in a rhombic dodecahedral simulation box under periodic boundary conditions, underwent one round of energy minimization using the conjugate gradient algorithm, and solvated using the TIP3P water model. Na^+^ and Cl^-^ ions were added to a final concentration of 50 mM and to neutralize the net system charge, followed by energy minimization via steepest descent algorithm. The system was then equilibrated in three steps: round one in the NVT ensemble for 5 ps at a integration step size of 0.5 fs, round two NpT ensemble for 12.5 ps (0.5 fs step size), and round 3 NpT ensemble for 25 ps (1 fs step size). Production runs were set to 298 K (v-rescale) and 1.013 bar with a 2 fs integration time step and coordinates were written every 5 ps. Short-range cutoffs for non-bonded interactions were set to 1.2 nm and long-range electrostatics were handled using the Particle Mesh Ewald method. Hydrogen bonds were converted to constraints using the LInear Constraint Solver. The force fields used in this work are listed in Fig. S5A, and available in the associated repository^65,66,91^. Performance differences of the different force fields are outlined in Fig. S5B-I. Bash scripts for simulation management and output post-processing as well as GROMACS simulation input files are available via the associated repository. Analyses were conducted using internal GROMACS command line tools and custom Python scripts using the MDAnalysis and MDtraj packages. The ContactCount to quantify binding of AT-hook peptides to the DNA was computed as outlined previously^92^. Briefly, distance-based contacts between hydrogen bond donors and acceptors of the core AT-hook residues and minor groove DNA bases were scored and combined into a collective variable. Initial structures of the D1 AT-hooks were obtained via the PEP-FOLD3 server, and termini were capped (N-terminal acetyl group and C-terminal amide group) using Pymol open source. Each AT-hook was manually placed next to B-form DNA at a distance of ≥ 1.5 nm using Pymol open source (Fig. S5H). The same simulation procedure was used as outlined above, and bound structures were identified via ContactCount and confirmed by visual inspection (Fig. S6). To estimate the relative binding strength of the D1 AT-hooks to DNA, we used the recently published steered MD simulation protocol^67^. Briefly, a pulling coordinate (*C*_*p*_) was defined by adapting ContactCount parameters, and the PLUMED interface was used to bias the simulation towards a reduction *C*_*p*_ over time, leading to unbinding of the AT-hook from the DNA, while the work invested in the unbinding was recorded. An example PLUMED input file specifying the simulation parameters can be accessed via the associated repository. We observed that one of the tested force fields lead to post-unbinding contributions to the work profile (Fig. S7B), leading to an exclusion of this force field from further simulations. At least 20 replicate simulations were run per AT-hook to determine the relative binding strength (Fig. S7F). There was no apparent difference in the binding strength based on the binding location of AT-hook #8 on the DNA (Fig. S6, Fig. S7C). To clean up noisy work profiles, we applied a minimum distance cutoff of 0.5 nm between the AT-hook core residues and the DNA, and disregarded work added to the system after this cutoff was reached (Fig. S7D). Lastly, the exponential average work profile of each AT-hook was bootstrapped to reduce outlier effects in the relative binding strength estimates (Fig. S7E, F). Bootstrapping was done by randomly sampling *k* trajectories (with replacement) 1000 times, where *k* is equal to the number of trajectories available for each AT-hook. The average work was calculated for each of the 1000 samples, and the mean and standard deviations of the average work where reported (Fig. S7F). Initial structures of DBM/NCR systems were extracted from the predicted structure of D1 from the AlphaFold Protein Structure Database (UniProt ID: P22058), and capped as described above. To simulate DNA binding, peptides were placed next to B-form DNA at a distance of ≥ 1.5, as before. To model DBM-NCR interactions, the peptides were placed in the same manner. Simulation protocols were employed as described above. DNA binding simulations were inspected after 200 ns of simulation time to determine promising conformations, which were continued for an additional 400 ns. Steered MD simulations were set up as described above to measure unbinding of DBM1 from DNA. Estimates of the enhanced individual binding strength of AT-hooks #1-3 as shown in Fig. 4F were determined by subsetting of the steered MD trajectories by order of unbinding. Visualizations were created using Pymol open source, and VMD. Termini were capped (N-terminal acetyl group and C-terminal amide group) using Pymol open source. Simulation prep, energy minimization, equilibration, and production runs were done using GROMACS 2024.1 with PLUMED 2.8.0 on the ETH Euler cluster. B-form DNA structure files were created using the NAB nucleic acid builder fd_helix command line tool by the Case group (https://casegroup.rutgers.edu/) accessible via the associated repository. Reference structures of AT-hooks bound to DNA were obtained from the protein databank (PDB) using accession numbers 2ezf, 2ezg, and 3uxw.

### DNA binding assays and in vitro phase separation

The D1 and mNG-D1 CDS were codon optimized for the expression in *Sf*9 insect cells and purchased from GenScript (Piscataway, NJ) with NheI and XhoI restriction sites at the 5’- and 3’-ends, respectively. The D1 and mNG-D1 gene were then cloned into pFB-MBP-WRN-10Xhis to generate pFB-MBP-D1co-10Xhis and pFB-MBP-mNG-D1co-10Xhis, respectively. The pFB-MBP-mNG-D1co-10Xhis construct was used as a template for cloning of the D1 variant constructs - the frameshift sequence found in *D*1^*m*04^ was pasted from genomic DNA using PCR. Proteins were expressed in *Sf*9 cells in SFX Insect serum-free medium (Hyclone) using the Bac-to-Bac expression system (Invitrogen), according to manufacturer’s recommendations. Frozen *Sf*9 pellets from 300 mL culture were resuspended in lysis buffer (50 mM Tris-HCl pH 7.5, 1 mM EDTA, 1:300 protease inhibitor cocktail [Sigma], 30 µg/ml leupeptin (Merck Millipore), 1 mM PMSF, 5 mM β-mercaptoethanol) and incubated at 4°C for 20 min. Glycerol was added to a final concentration of 25%, NaCl was added to a final concentration of 305 mM and the solution was incubated at 4°C for 30 min. The cell suspension was centrifuged at 55,000 g at 4°C for 30 min. The soluble extract was incubated with amylose resin (New England Biolabs) at 4°C for 1 h. The resin was washed with amylose wash buffer I (50 mM Tris-HCl pH 7.5, 5 mM β-mercaptoethanol, 1 M NaCl, 10% glycerol, 1 mM PMSF), followed by one wash with amylose wash buffer II (50 mM Tris-HCl pH 7.5, 5 mM β-mercaptoethanol, 300 mM NaCl, 10% glycerol, 1 mM PMSF). Proteins were eluted using amylose elution buffer (50 mM Tris-HCl pH 7.5, 5 mM β-mercaptoethanol, 300 mM NaCl, 10% glycerol, 1 mM PMSF, 10 mM maltose [Sigma]). The MBP-tagged variants were incubated with PreScission protease (3.3 mg PreScission protease per 100 mg of tagged protein) at 4°C for 20 h to cleave the MBP-tag. Subsequently, imidazole was added to a final concentration of 10 mM and the solution was incubated with pre-equilibrated Ni-NTA agarose resin (Qiagen) at 4°C for 1 h, in agitation. The resin was washed with Ni-NTA buffer I (50 mM Tris-HCl pH 7.5, 5 mM β-mercaptoethanol, 1 M NaCl, 10% glycerol, 1 mM PMSF, 30 mM imidazole) and subsequently with Ni-NTA buffer II (50 mM Tris-HCl pH 7.5, 5 mM β-mercaptoethanol, 150 mM NaCl, 10% glycerol, 1 mM PMSF, 30 mM imidazole). Proteins were eluted with Ni-NTA elution buffer (50 mM Tris-HCl pH 7.5, 5 mM β-mercaptoethanol, 100 mM NaCl, 10% glycerol, 1 mM PMSF, 300 mM imidazole) and dialyzed against 1 L of dialysis buffer (50 mM Tris-HCl pH 7.5, 5 mM β-mercaptoethanol, 100 mM NaCl, 10% glycerol, 1 mM PMSF) at 4°C for 1 h. Fractions containing high protein concentration as estimated by Bradford were pooled, aliquoted, snap-frozen in liquid nitrogen and stored at −80°C. The oligonucleotide-based 62 bp-long dsDNA substrates were prepared from single-stranded oligonucleotides (forward) and single-stranded reverse complements (reverse). The forward oligonucleotides were ^32^P-labeled at the 3’ terminus with [α-^32^P]dCTP (Hartmann Analytic) and terminal transferase (New England Biolabs) according to the manufacturer’s instructions. Free nucleotides were removed with Micro Bio-Spin P-30 Tris chromatography columns (Biorad). Cy-3 labelled plasmid DNA for *in vitro* phase separation experiments were produced from pMK-RQ_192xAATAT (pMK-RQ backbone with an inserted tract containing roughly 192 repeats of AATAT) using the Label IT^®^ Nucleic Acid Labeling Kit (MIR 3625, mirus bio) according to the manufacturer’s instructions. Electrophoretic mobility shift assay (EMSA) experiments (15 µl volume) were performed in binding buffer containing 25 mM Tris-acetate pH 7.5, 1 mM DTT, 3 mM MgCl_2_, 0.1 mg/ml BSA (New England Biolabs) and 1 nM DNA substrate (in molecules). After the addition of D1 protein, the reactions were incubated on ice for 15 min. Loading dye (50% glycerol, bromophenol blue) was added and the products were separated by 4% polyacrylamide (19:1 acrylamide-bisacrylamide, Bio-Rad) native gel electrophoresis in Tris-Acetate-EDTA (TAE) buffer. The gels were dried on a 17 CHR paper (Whatman) and exposed to storage phosphor screen (GE Healthcare) and scanned by a Typhoon Phosphor Imager (FLA 9500, GE Healthcare). Band intensities were analyzed using ImageJ/FIJI. Phase separation assays were done in 384 well microplates (Greiner Item-No. 781096, Greiner Bio-One) pre-treated for >1 h with 10 mg/mL BSA (A1391, PanReac AppliChem) in ddH2O, rinsed 3x with ddH2O and air-dried. Reactants were mixed to their final concentrations directly in the wells of the BSA-coated 384 well microplates, incubated for 15 minutes at room temperature, and spun down for 1 minute at 10 RCF. Images for quantification were recorded using a Nikon Ti2-E widefield microscope (40x, air) with an automated XY-stage. Image acquisition comprised 3 images per well; all experiments were done in duplicates. Widefield images were analysed via a custom CellProfiler (version 4.2.6) pipeline (accessible via the online repository). High-resolution confocal images for visualization were recorded on a Leica TCS SP8 inverted confocal microscope (64x, oil).

### Mouse cell culture and image analysis

Chromocenter clustering assays in cultured mouse cells were conducted as described previously^70^. Briefly, NIH-3T3 cells were cultured in high glucose DMEM with pyruvate supplemented with 10% FBS (Life Technologies Limited, UK). D1 and D1 variant sequences were amplified via PCR from genomic fly DNA and cloned into a pCDNA3 vector downstream of a mNeonGreen (mNG) fluorescent tag. Cells were grown for 24 hours on round 12 mm coverslips to 50% confluency, followed by transfection with Lipofectamine 2000 Transfection Reagent (Invitrogen, USA), according to manufacturer’s instructions. Cells were fixed in 3.6% formaldehyde 48h post-transfection and permeabilized for 5 minutes with 1x PBS containing 0.4% Triton-X 100. Cells were mounted with VECTASHIELD^®^ Antifade Mounting Medium with DAPI (Vector Laboratories, USA). The mounting medium was additionally supplemented with 1:1000 Sphero Fluorescent Yellow Particles 0.5 µm (Spherotech, USA) to enable estimation of expression levels across samples. Imaging of nuclei and fluorescent beads was done on a Leica SP8 AOBS laser scanning confocal microscope using a 63x oil immersion objective (NA = 1.4). Experiments consisted of 3-4 independent replicate samples; 4 images of fluorescent beads in different regions of the sample were recorded per sample. Nuclei and chromocenters were segmented on the DAPI channel (excitation at 405 nm), and fluorescent beads were segmented on the mNG channel (excitation at 488 nm) using CellProfiler (version 4.2.6), and fluorescent intensities were measured within the segmented objects. Normalized concentrations were determined by normalization of fluorescent intensity to the median upper-quantile intensity of fluorescent beads. Image data was manually filtered to exclude wrongly segmented nuclei, any nuclei above an empirically determined size threshold, as well as nuclei containing more than 65 chromocenters. Data processing and statistical analysis and visualization were done in Python and R. Representative microscopy images were prepared with ImageJ/FIJI. For live imaging, NIH-3T3 cells were cultured in an 8-well ibiTreat imaging chamber slide (ibidi GmbH, Germany) in high glucose DMEM with pyruvate supplemented with 10% FBS (Life Technologies Limited, UK). Transfection reactions were carried out as described above, using Lipofectamine 2000 Transfection Reagent (Invitrogen, USA). Imaging commenced 24 hours post transfection using a Visitron Spinning Disc confocal microscope. Image data was analyzed with ImageJ/FIJI.

### RNA-Sequencing and analysis

For each sample, 40 adult testes from 0-1 day old males were dissected into RNAse-free 1x PBS and flash frozen in liquid nitrogen. Total RNA was extracted using the RNeasy® Mini Kit 50 (Qiagen). The samples were post-processed using the Relia Prep RNA Clean Up & Concentration System kit (Promega). RNA quality was controlled using a Nanodrop and a Qubit RNA analyzer. Samples of sufficient quality (RIN > 9) were subjected to library preparation (Illumina TruSeq mRNA kit) followed by sequencing using Illumina NovaSeq 6000 (single-read, 100 bp) at the Functional Genomics Center Zürich (FGCZ). The resulting raw reads were cleaned by removing adaptor sequences, low-quality end trimming, and removal of low-quality reads using BBTools v38.18 (https://sourceforge.net/projects/bbmap). The exact commands used for quality control are available on the Methods in Microbiomics webpage (https://methods-in-microbiomics.readthedocs.io/en/latest/preprocessing/preprocessing.html). Transcript abundances were quantified using Salmon v1.10.1 and BDGP6.32. Differential gene expression analysis was performed using the Bioconductor R package DESeq2 v1.37.4.

### Electron microscopy

Testes were dissected in freshly prepared fixation solution (2.5 % glutaraldehyde in 0.1 M sodium cacodylate buffer) and stored at 4 °C until further processing. TEM was performed at the Center for Microscopy and Image Analysis (University of Zurich, Switzerland). Image analysis was performed using Maps Viewer, color overlays were created in Photoshop (Adobe).

## QUANTIFICATION AND STATISTICAL ANALYSIS

Data processing was achieved via custom Python scripts using NumPy and Pandas, or using the dplyr package in R. Statistical tests were conducted in R, using wilcox.test from stats. All Mann-Whitney U tests were done assuming two-sided hypotheses. See the respective sections above and the online repository at https://gitlab.ethz.ch/bfruehbauer/chromocenter_nucmech2025 for in-depth information on specific analyses.

