## Supplementary Tables and Figures for "Nuclear mechanostability emerges from satellite DNA condensation into chromocenters"

**SUPPLEMENTAL INFORMATION**

**Supplementary Table 1. D1 gRNA sequences and combinations used to generate D1 variants.**

| gRNA sequence | Cleavage site | Duplex group (ii) | Cassette (iii) |
|---|---|---|---|
| ACTGGAGCCCTTGTTCTTGG | K41 | A | 1 |
| GTCCTCCTCCCCCTCGTCTT | D69 | B | 1 |
| AGTACTGGCTACGTACCCAC | P216 | D | 1 |
| TCGTCCGCGCGGTCGTCCAA | P224 | C, F | 1 |
| CTTGCGCTTCTTGGCGGAAC | S118 | C, E | 2 |
| CTGTCGAGTCCAATGGCGAT | G188 | A, E | 2 |
| CGAGCTGTGTTCCTCCTCCC | D248 | F | 2 |
| GGGCCAACACAACTCCAAGA | K308 | B, D | 2 |

**Video S1. Loss of D1 leads to nuclear deformation and MN formation.** Z-stack videos of fixed testes from control ($D1^{KO/+}$, left) and D1 mutant ($D1^{KO/LL}$, right) 0 – 1 day old adult flies stained for Vasa (grey), Lamin B (green) and DAPI (blue). Yellow arrowheads indicate examples of strong nuclear deformations. Magenta arrowheads mark MN. Scale bars are 10 µm.

**Video S2. Adult spermatogonia experience more mechanical forces than their larval counterparts.** Synchronized time-lapse videos of living larval (left) and adult (right) testes expressing Aub-GFP to visualize the cytoplasm of germ cells. Note the peristaltic contractions in the adult testes caused by a muscle sheath.

### SUPPLEMENTARY FIGURES

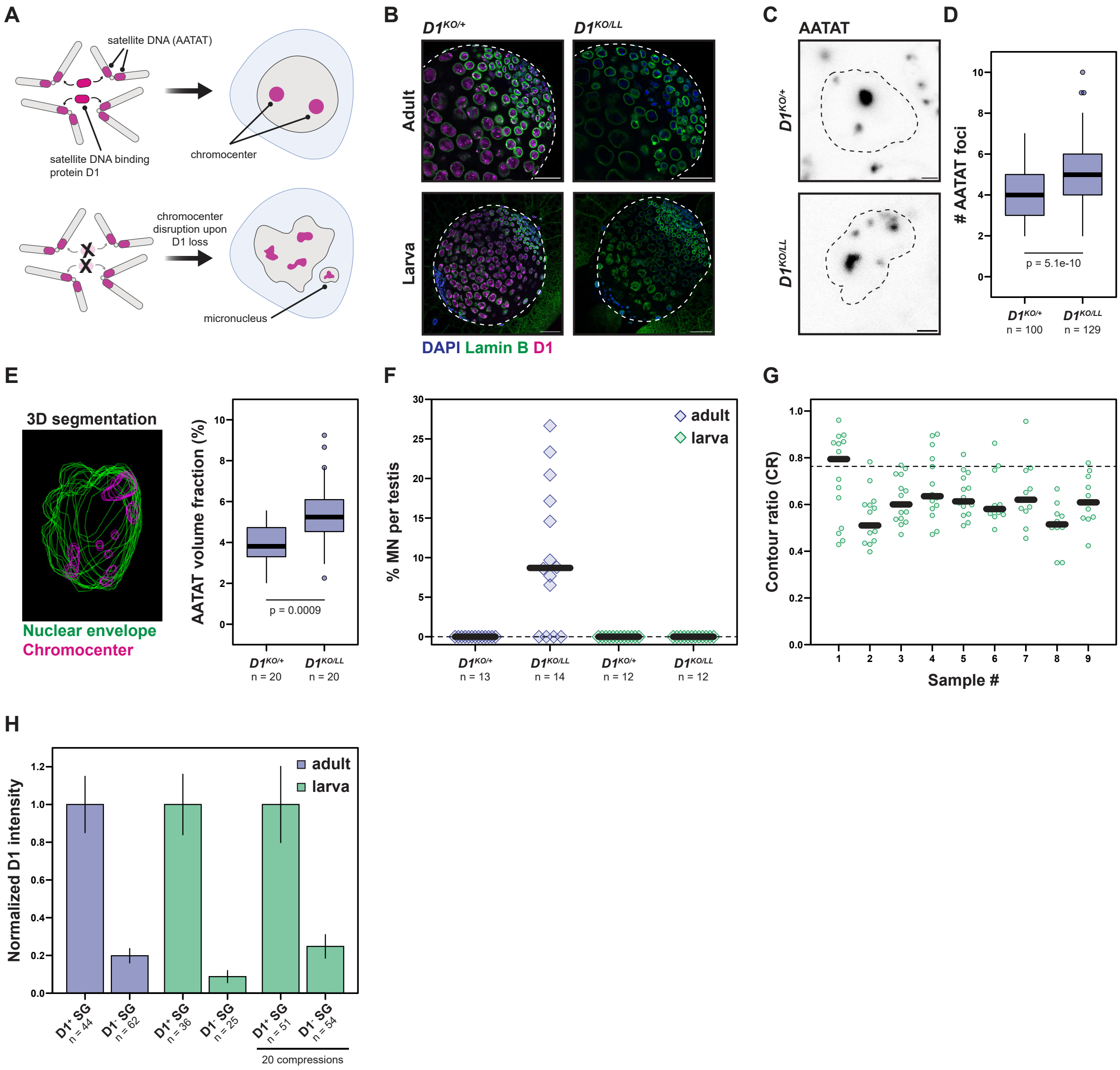


**Figure S1. Loss of D1 causes chromocenter disruption, nuclear deformation and MN formation in adult and mechanically challenged larval spermatogonia** (A) Schematic of D1 function in *Drosophila* male germ cells. (B) Control (*D1*$^{KO/+}$) and *D1* mutant (*D1*$^{KO/LL}$) adult and larval testes stained for D1 (magenta), Lamin B (green) and DAPI (blue). Outlines mark tissue boundaries. Scale bars for adult and larval tissue are 25 µm and 50 µm, respectively. (C) FISH against the AATAT satellite DNA repeat in control and *D1* mutant adult spermatogonia. Maximum intensity projections are shown, and outlines indicate nuclear boundaries. Scale bars: 1 µm. (D) Number of AATAT foci in control and *D1* mutant adult spermatogonia. (E) Example of 3D segmentation of a nucleus (green) and its chromocenters (magenta) and quantification of AATAT volume fraction in nuclei from control and D1 mutant adult spermatogonia. (F) % MN per testis from the indicated genotypes and developmental stages. (G) CR values for compressed *D1* mutant larval spermatogonia (from Fig. 1G) grouped by cells from the same testes. (H) Normalized D1 expression levels in D1-positive and D1-negative spermatogonia of adult, larval and compressed larval testes. Indicated p values are from Mann-Whitney U-tests. For the CR and MN plots, the thick black line indicates the median while the dashed line represents the median CR (0.76) in adult *D1* heterozygous spermatogonia.

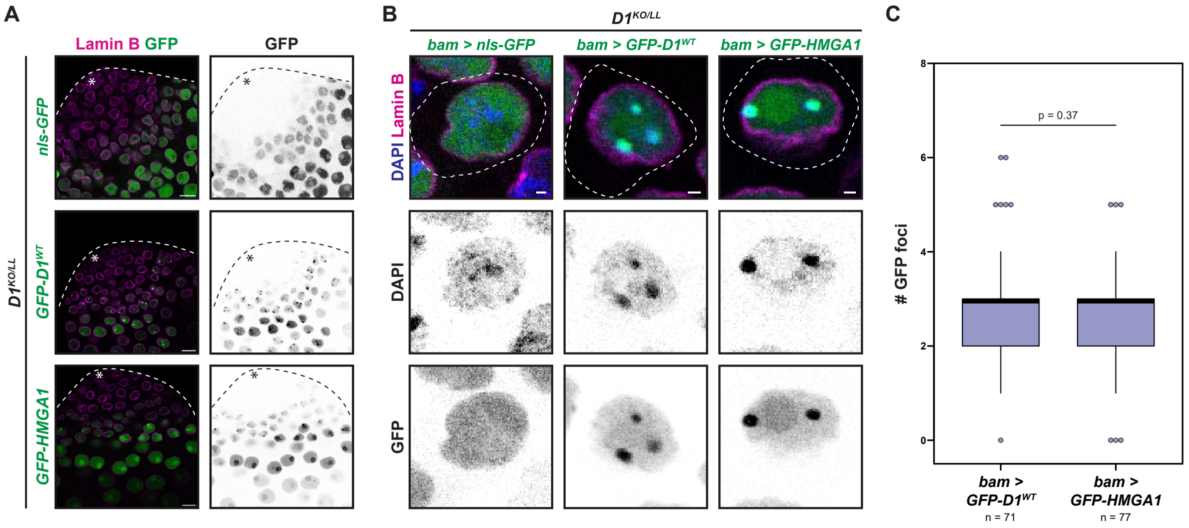


**Figure S2. HMGA1 restores chromocenters in *D1* mutant spermatogonia.** (A) Overview of *bam-Gal4* driven expression of the indicated GFP-tagged constructs (green) in *D1*$^{KO/LL}$ testes stained for Lamin B (magenta). Asterisks indicate the positions of the stem cell niche. (B) *D1* mutant spermatogonia expressing NLS-GFP, GFP-D1$^{WT}$ and GFP-HMGA1 co-stained for DAPI (blue) and Lamin B (magenta). Scale bars: 1 µm. (C) Quantification of GFP foci per nucleus from (B). Indicated p value is from a Mann-Whitney U-test.

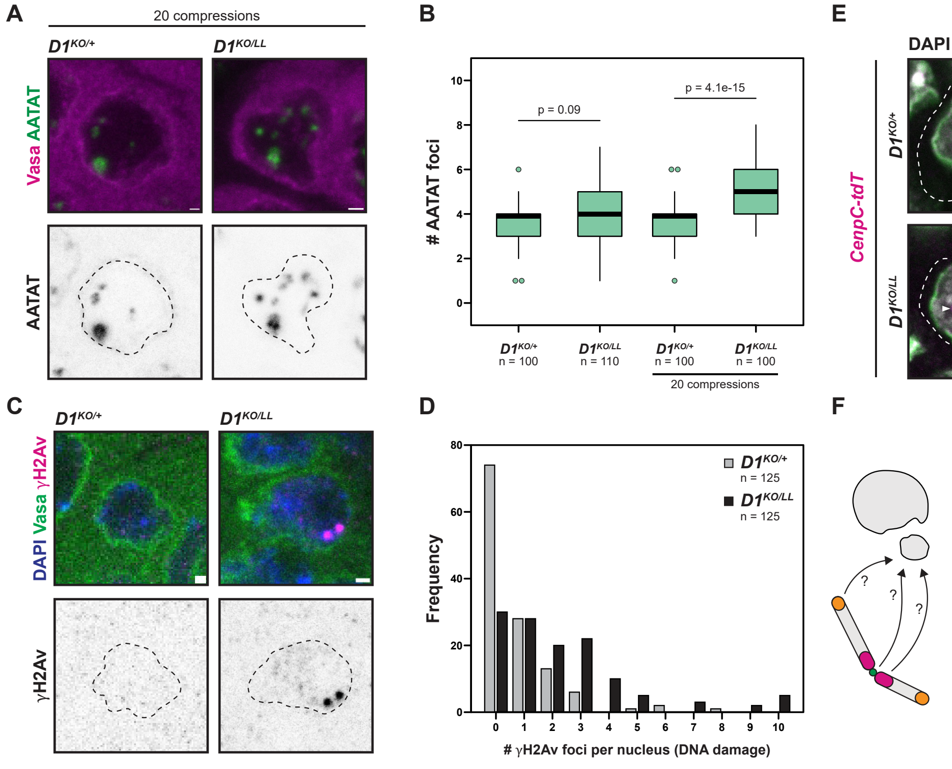


| Probe | Chr. | n | % in MN |
|---|---|---|---|
| Telomeres (HOAP) | all | 38 | 78.9 |
| Centromeres (CenpC) | all | 48 | 39.6 |
| AATAACATAG | 2, 3 | 34 | 17.6 |
| AATAT | X, Y, 3, 4 | 20 | 35 |
| AATAC | Y | 37 | 18.9 |
| 359bp | X | 30 | 13.3 |
| AACAC | 2, Y | 33 | 15.2 |
| Dodeca | 3 | 49 | 10.2 |
| AAGAT | 4 | 17 | 5.9 |
| AATAGAC | Y | 20 | 0 |
| TAGA | X | 14 | 0 |

**Figure S3. Interphase micronuclei in *D1* mutant spermatogonia contain fragmented chromosomes.** (A) FISH against the AATAT satellite (green) in control and *D1* mutant larval spermatogonia after 20 compressions co-stained with Vasa (magenta). Outlines indicate nuclear boundaries. (B) Quantification of number of AATAT foci per nucleus in control and *D1* mutant larval testes before and after 20 compressions. Indicated p values are from Mann-Whitney U-tests. (C) Adult spermatogonia from the indicated genotypes stained for γH2Av (magenta), Vasa (green) and DAPI (blue). Outlines indicate nuclear boundaries. (D) Quantification of γH2Av-foci per adult spermatogonia from control and *D1* mutant testes. (E) Control and *D1* mutant spermatogonia expressing CenpC-tdT (magenta) stained for Lamin B (green) and DAPI (gray). Centromeres are marked with arrowheads. Outlines indicate cellular boundaries. (F) Quantification of MN contents using fluorescently tagged proteins labeling telomeres (HOAP-GFP), centromeres (CenpC-tdT), antibody staining (Prod) and FISH against the indicated satellite DNA loci. The final bin contains all nuclei containing 10 or more foci. All scale bars are 1 μm.

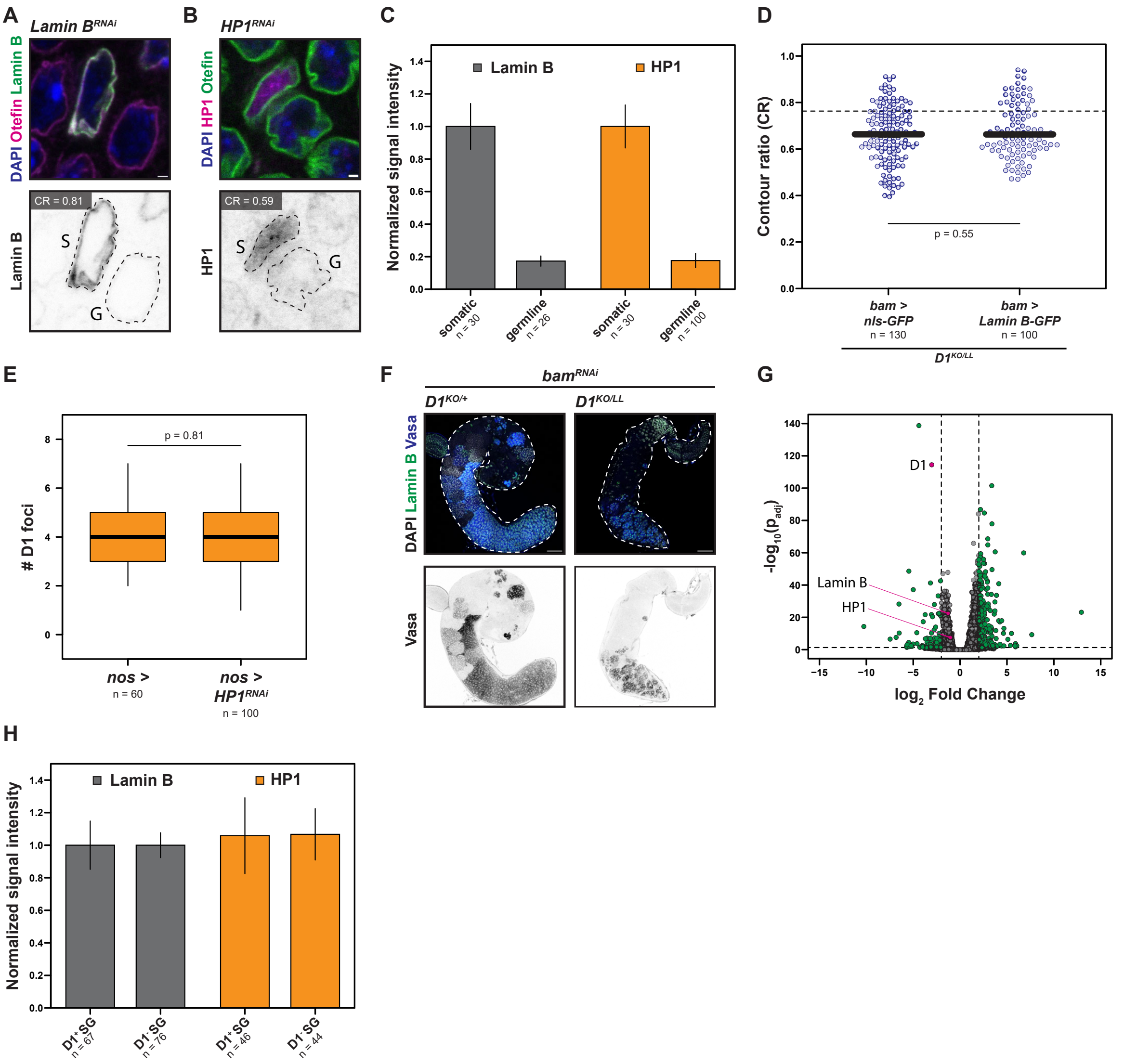


**Figure S4. Lamin B and HP1 levels are not significantly altered in *D1* mutant spermatogonia.** (A, B) Representation of a somatic 'S' and a germ cell 'G' in an adult testis expressing *Lamin B*$^{RNAi}$ (A) and *HP1*$^{RNAi}$ (B) under the control of *nos-gal4* stained for Lamin B/Otefin (green), Otefin/HP1 (magenta) and DAPI (blue). Outlines represent nuclear boundaries. CR value corresponds to the represented germ cell. Scale bar 1 µm. (C) Knockdown efficiencies of Lamin B and HP1 respectively, normalized to the mean intensity value of each category. (D) CR of adult spermatogonia from the indicated genotypes. *bam > NLS-GFP* datapoints are replotted from Fig. 1N. (E) Number of D1 foci in control (*nos >*) and HP1-depleted adult spermatogonia. (F) Control and *D1* mutant adult testes enriched for spermatogonia using *nos>bam*$^{RNAi}$ stained for Lamin B (green), Vasa (blue) and DAPI (grey). Scale bars: 50 µm. (G) Volcano plot of differentially expressed genes in control vs. *D1* mutant testes in a *nos>bam*$^{RNAi}$ background. Genes passing a cutoff of $\log_2FC>2$, $p_{adj}<0.05$ are highlighted in green. D1, Lamin B and HP1 are labeled in magenta. (H) Intensities of Lamin B and HP1 in $D1^{+}$ versus $D1^{-}$ spermatogonia in mosaic testes normalized to the expression in somatic cells.

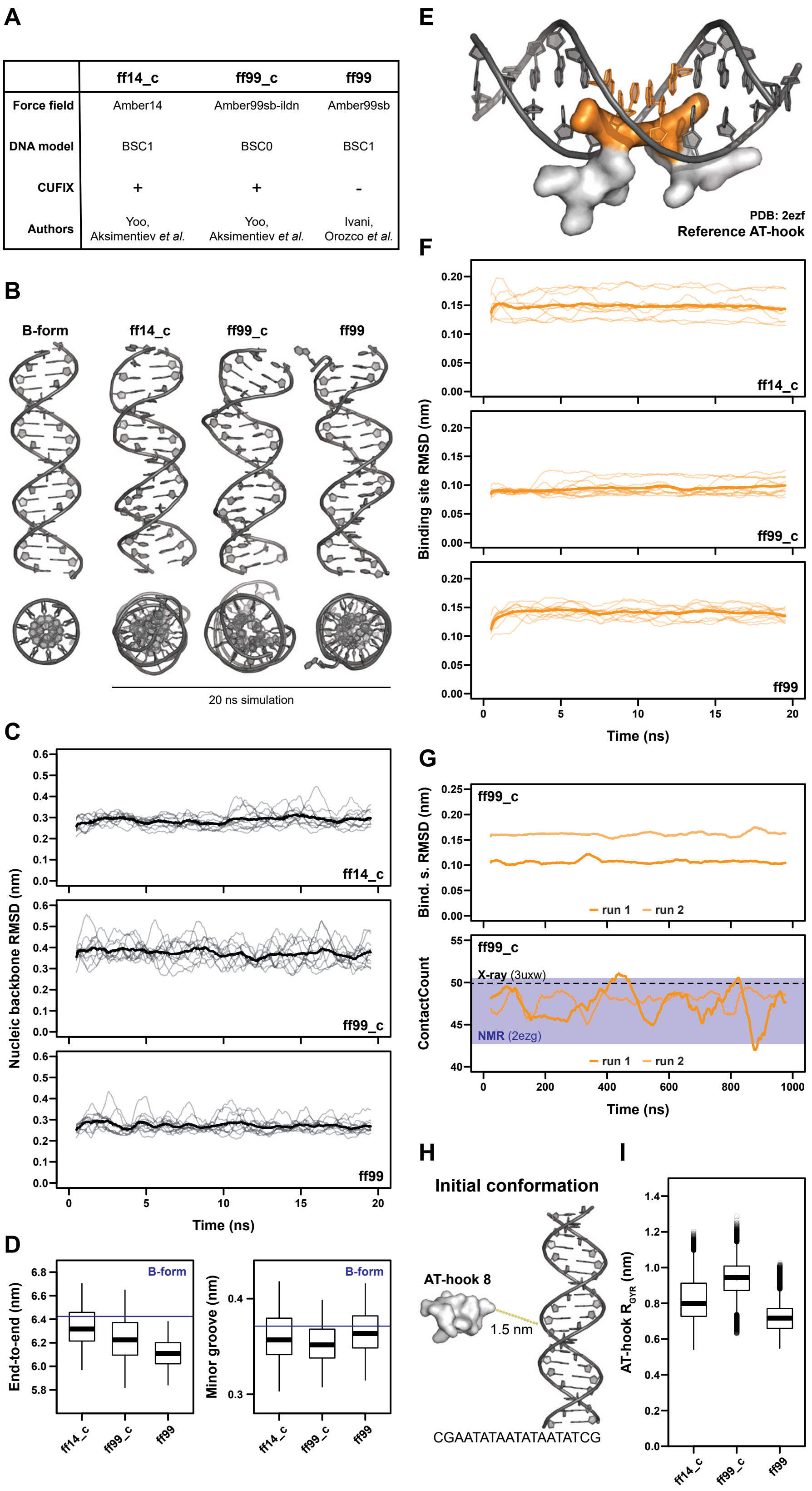


**Figure S5. Amber99sb-ildn bsc0 cufix recapitulates AT-hook DNA interactions.** (A) Overview of force fields considered in this work. (B) Side view (top) and top view (bottom) of B-form DNA and DNA conformations obtained from MD simulation. (C) Structural deviation trajectories of the DNA backbone in the different force fields. Thick lines show the average over the replicate trajectories. (D) Conformational distributions of DNA end-to-end distances (left) and the distance of the nitrogen and oxygen atoms in a minor-groove hydrogen bond (right). The blue line indicates values measured in the initial B-form DNA conformation. (E) Reference structure of an AT-hook bound to DNA (PDB: 2ezf). (F) Positions highlighted in orange from (E)

are used to investigate how well the force fields can reproduce the experimentally determined structure in terms of the RMSD over time. (G) Stability of the reference AT-hook structure in µs simulations. RMSD over time (top) and ContactCount over time (bottom) are shown. Reference ContactCounts derived from a crystal structure (PDB: 3uxw) and an NMR ensemble (PDB: 2ezg) are indicated. (H) Example conformation for D1 AT-hook DNA binding prediction. The peptide is kept at a distance larger than the short-range cutoff from the DNA to avoid initial sampling bias. (I) Conformational distribution of D1 AT-hooks (pooled over AT-hooks and replicate simulations) in terms of the peptides' radius of gyration ($R_{GYR}$).

Bound conformations

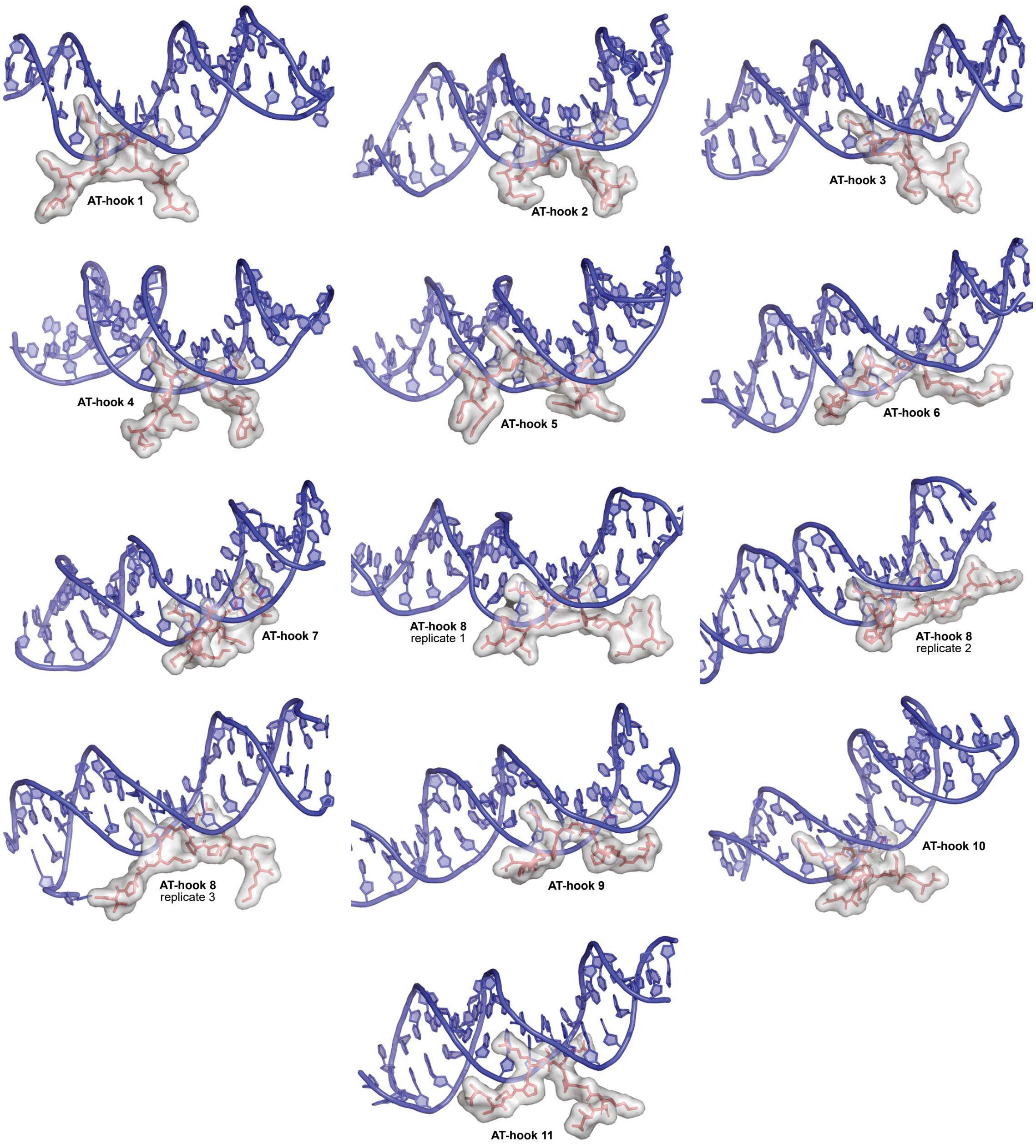


**Figure S6. Bound conformations of D1 AT-hooks.** Representative structures of all 11 D1 AT-hooks bound to 3xAATAT.

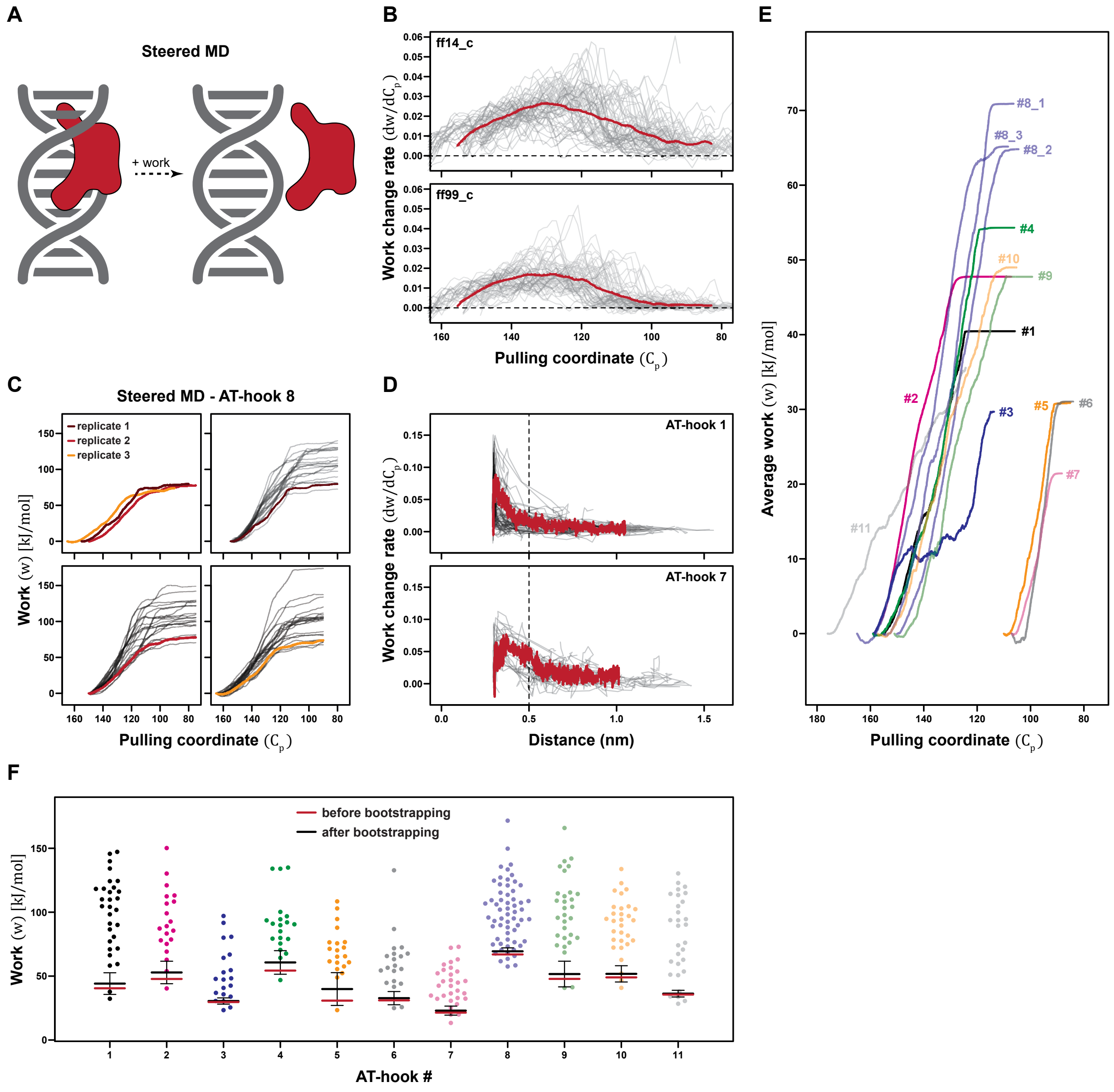


**Figure S7. Steered MD simulations reveal sequence-dependent differences in D1 AT-hook binding strength.** (A) Schematic of steered MD simulation strategy. (B) Estimate of post-unbinding contributions to work measured in steered MD through numerical derivation of work as a function of the pulling coordinate ($C_p$). Red lines indicate averages over all trajectories. (C) Binding site influence on estimated unbinding work shown by using differentially bound conformations of AT-hook 8; Boltzmann-weighted averages and individual work are shown for the 3 replicates. (D) AT-hook to DNA distance-dependent contribution to overall work in steered MD simulations. AT-hooks 1 and 7 are shown as examples. The cutoff-distance of 0.5 nm that was used in work-profile post-processing is indicated. Red lines indicate averages over all trajectories. (E) Boltzmann-weighted average work curves of the individual AT-hooks after distance-dependent correction of the work profiles. (F) Work for unbinding as measured for each AT hook. Red line segments indicate the raw Boltzmann-weighted average, black line segments show the corrected Boltzmann-weighted average after bootstrapping.

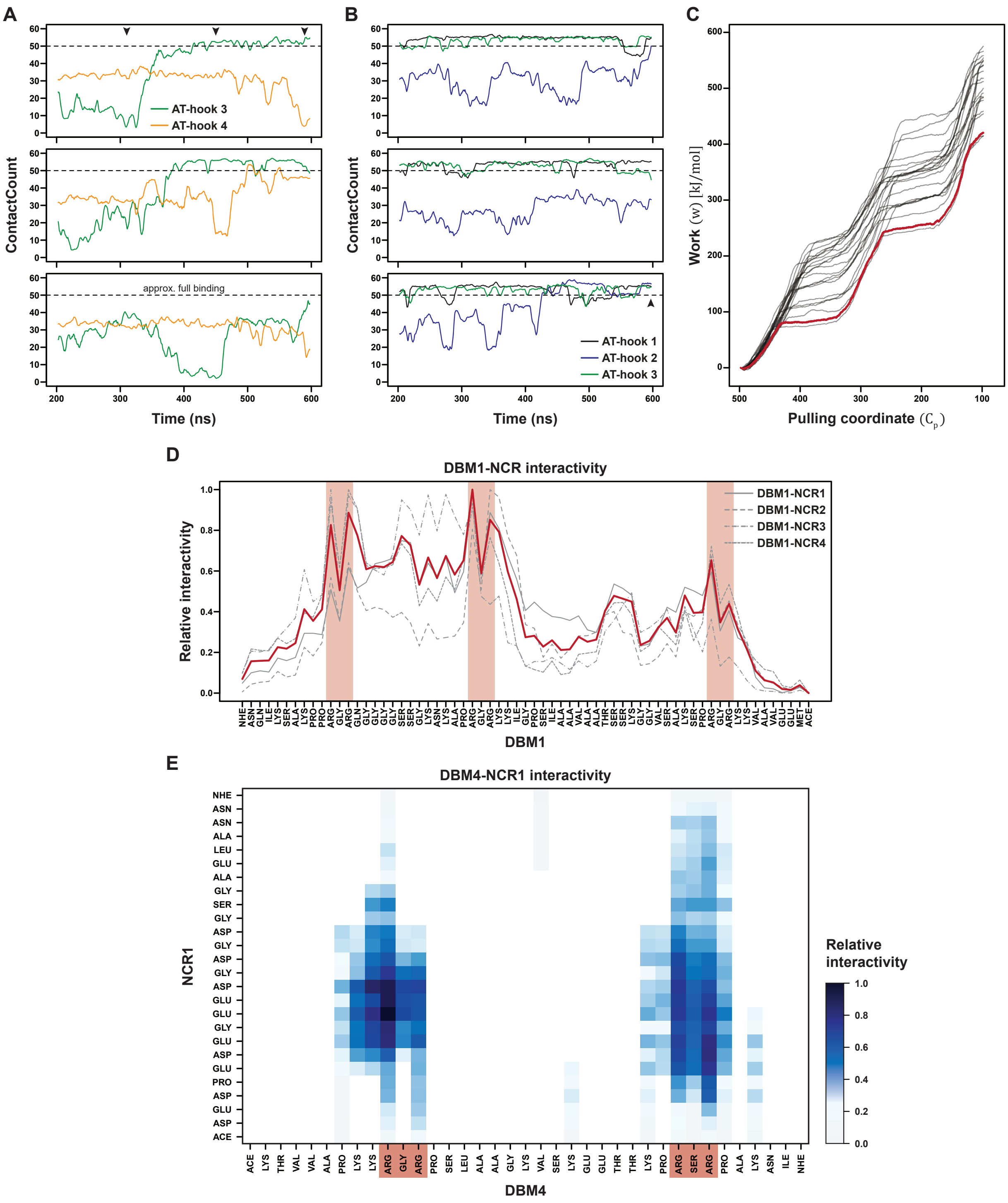


**Figure S8. Cooperative DBM-DNA interactions are disrupted by charge-based DBM-NCR interactions.** (A) ContactCount for individual AT-hooks in simulations of ATH3-NCR1-ATH4 over time. The dashed line indicates the average ContactCount of a fully bound AT-hook. Arrowheads correspond to conformations shown in Fig. 4D. (B) ContactCount for individual AT-hooks in simulations of DBM1 over time. The dashed line indicates the average ContactCount of a fully bound AT-hook. Arrowhead corresponds to conformations shown in Fig. 4E. (C) Work profile for DBM1 derived from steered MD simulation. Red line shows the Boltzmann-weighted average. (D) 2D interaction profile of DBM1 with NCRs. The red line shows the average over the different DBM1-NCR systems. AT-hook locations are highlighted in light red. (E) Per-residue interaction map of DBM4 and NCR1. AT-hook locations are highlighted in light red.

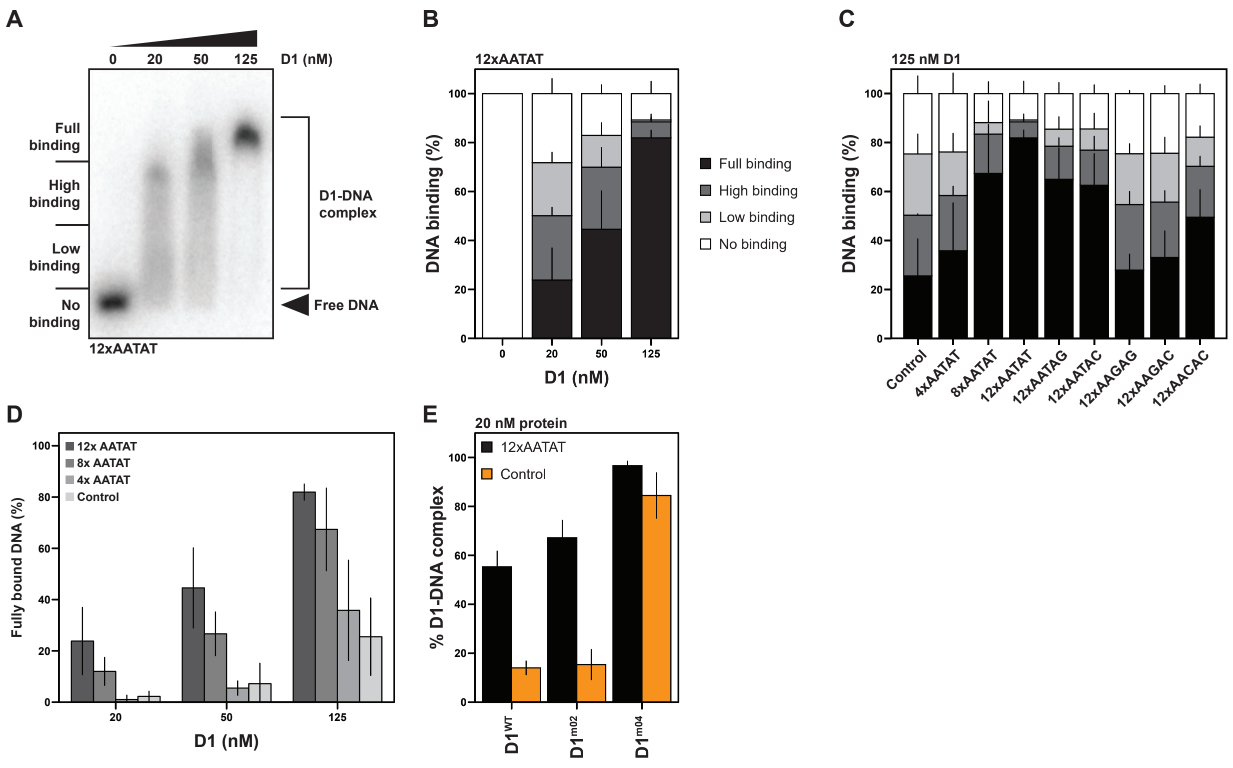


**Figure S9. D1 shows preferential binding to AATAT oligonucleotides.** (A, B) Representation (A) and quantification (B) of electrophoretic mobility shift assay (EMSA), showing changes in 12xAATAT DNA migration at increasing D1 concentrations. (C) Quantification of D1 binding to the indicated oligonucleotides at a protein concentration of 125 nM. (D) Quantification of D1 binding to the indicated oligonucleotides at increasing concentrations. (E) Quantification of D1 wildtype and D1 variant binding to control and 12xAATAT oligonucleotides. Barplots show means and standard deviations.

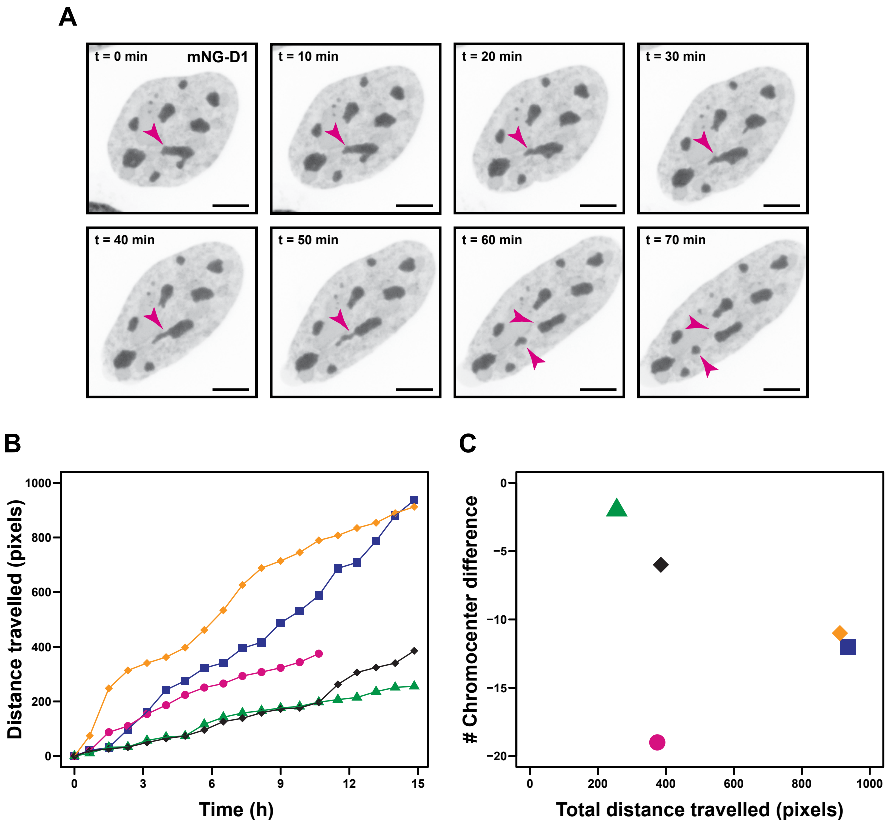


**Figure S10. D1-dependent chromocenter clustering is associated with cell migration.** (A) A chromocenter in a NIH-3T3 nucleus (arrowhead) undergoes fission upon nuclear stretching on an hour-long timescale. Scale bars: 5 µm. (B) Cumulative migrated distance of nuclei corresponding to Fig. 5C. (C) Change in the number of chromocenters over the course of 15 hours in relation to the total migration distance of the cells corresponding to Fig. 5C.

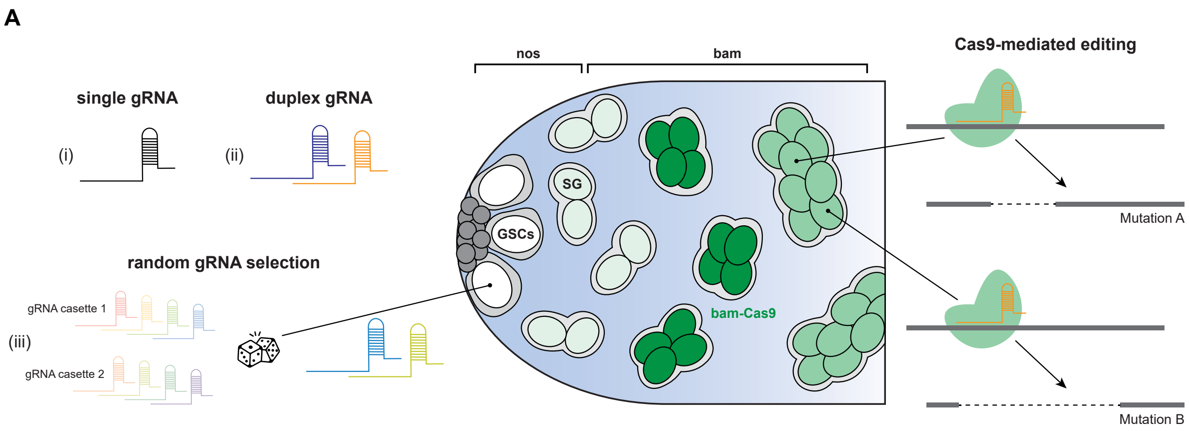


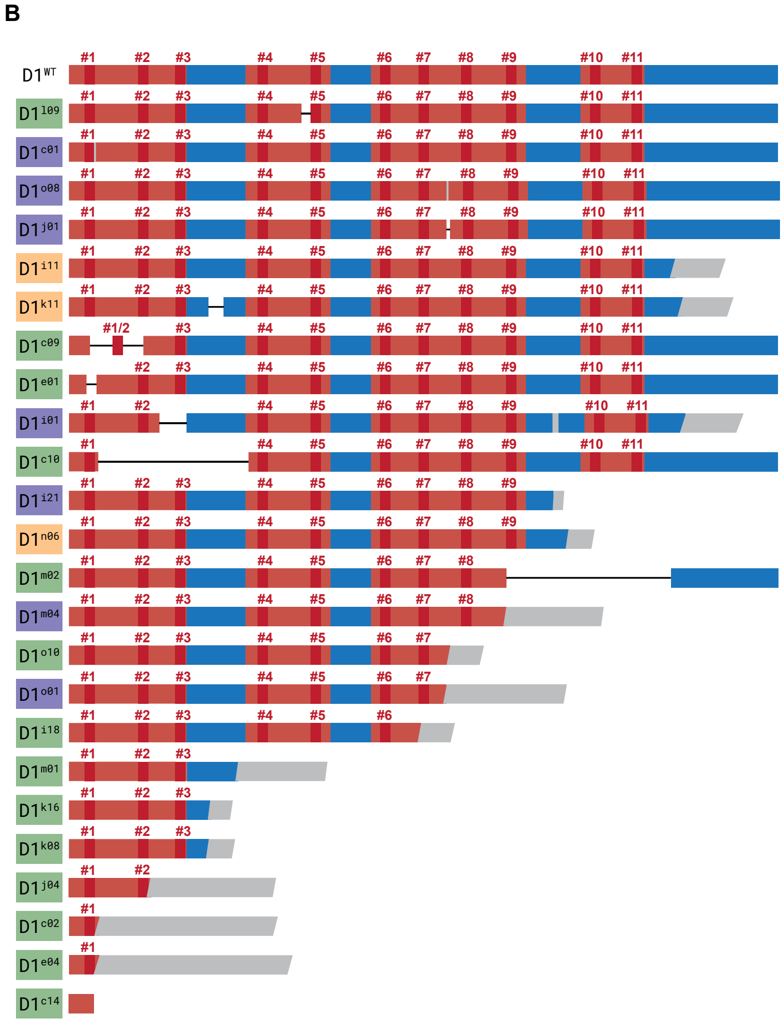


| Variant | ATHs | Charge | Clust. sc. | Partitioning |
|---|---|---|---|---|
| D1WT | 11 | -1 | 9.78 | 1.76 |
| D1l09 | 11 | -5 | 10.90 | 1.64 |
| D1c01 | 11 | -2 | 15.31 | 1.80 |
| D1o08 | 11 | -1 | 12.34 | 1.86 |
| D1j01 | 11 | 0 | 11.56 | 1.82 |
| D1i11 | 11 | +10 | 33.27 | 2.22 |
| D1k11 | 11 | +17 | 37.07 | 2.39 |
| D1c09 | 10 | -7 | 12.94 | 1.56 |
| D1e01 | 10 | -4 | 8.94 | 1.71 |
| D1i01 | 10 | +8 | 26.00 | 1.78 |
| D1c10 | 9 | -1 | 10.52 | 1.55 |
| D1i21 | 9 | +8 | 20.72 | 1.78 |
| D1n06 | 9 | +14 | 26.94 | 2.18 |
| D1m02 | 8 | +3 | 16.77 | 1.42 |
| D1m04 | 8 | +24 | 33.37 | 1.82 |
| D1o10 | 7 | +7 | 18.51 | 1.33 |
| D1o01 | 7 | +25 | 34.24 | 1.56 |
| D1i18 | 6 | +6 | 18.91 | 1.27 |
| D1m01 | 3 | +8 | 18.31 | 1.23 |
| D1k16 | 3 | +15 | 25.24 | 1.18 |
| D1k08 | 3 | +15 | 29.67 | 1.08 |
| D1j04 | 2 | +19 | 22.07 | 1.17 |
| D1c02 | 1 | +19 | 22.16 | 1.10 |
| D1e04 | 1 | +21 | 29.65 | 1.05 |
| D1c14 | 0 | +3 | 25.15 | 0.92 |

**Figure S11. Establishing a *D1* mutant library.** (A) Schematic representation of the mutagenesis approaches used in the present study. D1 was targeted by (i) a ubiquitously expressed single gRNA (BDSC:84068), (ii) ubiquitously expressed gRNA duplexes, or (iii) gRNA duplexes randomly selected from two gRNA cassettes in germline stem cells (GSCs) by *nos*-driven expression of phiC31 (see Materials and Methods for further details). Cas9 was expressed under the control of the *bam* promoter to induce non-clonal mutations in spermatogonia. (B) Selected D1 variant proteins screened for their chromocenter clustering properties in mouse cells. Shown are their number of AT-hooks, net charge, clustering scores and partitioning into chromocenters. Color coding of variants corresponds to Fig. 5G.

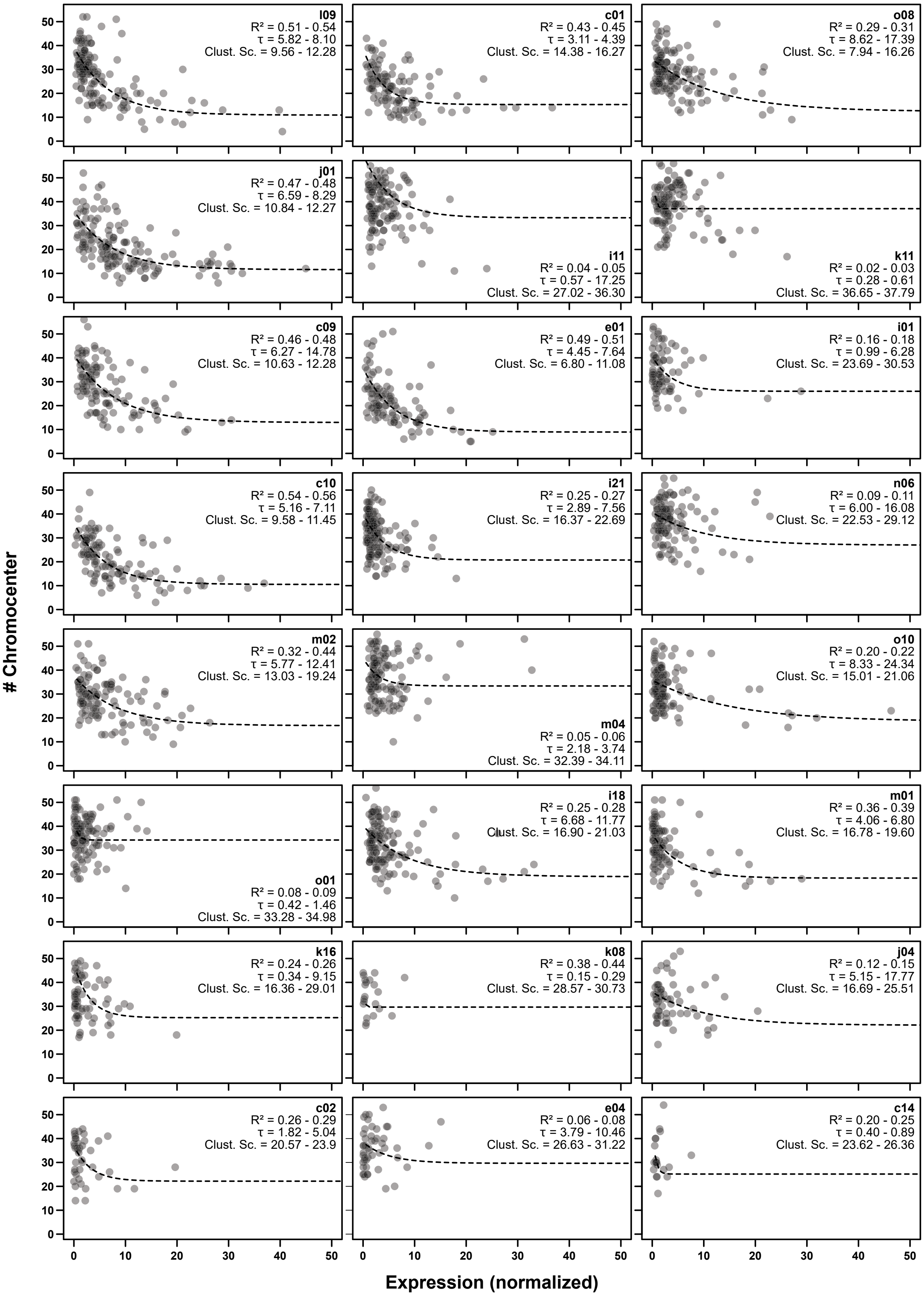


**Figure S12. Chromocenter clustering by D1 variant proteins.** Number of chromocenters per nucleus in relation to normalized expression of D1 variant proteins 48h post transfection. The ranges for $R^2$, time constant (τ) and clustering score are shown for each variant following bootstrapping.

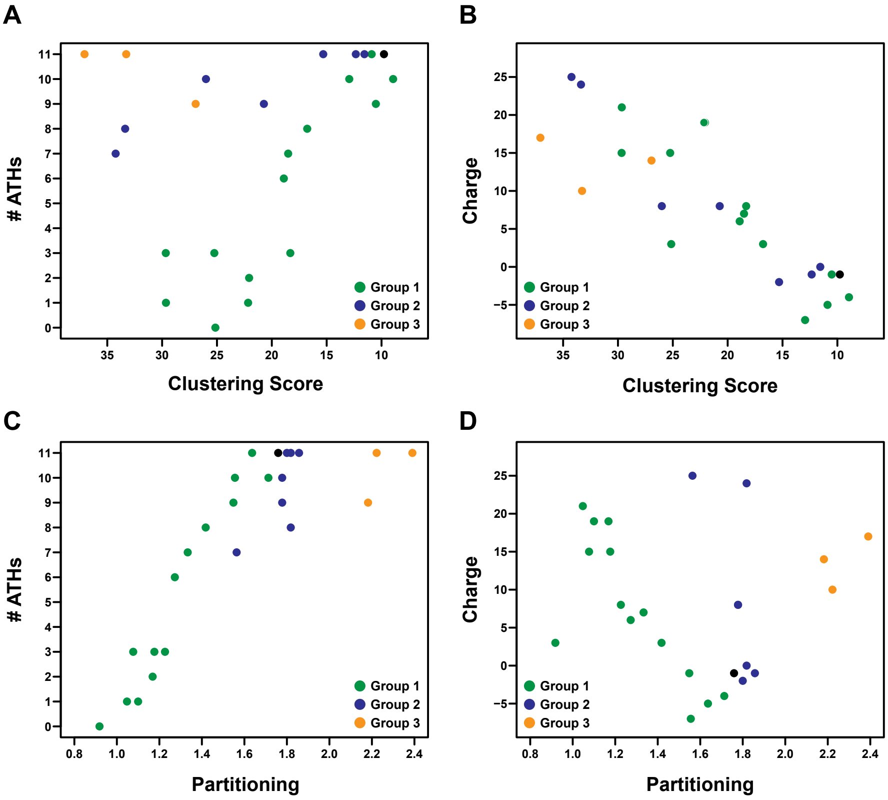


**Figure S13. D1-dependent satellite DNA clustering is regulated by number of AT-hooks and net charge**. (A-D) Relationship between AT-hook number (A, C) and net charge (B, D) on clustering score (A, B) and partitioning (C, D) in mouse NIH-3T3 fibroblasts.

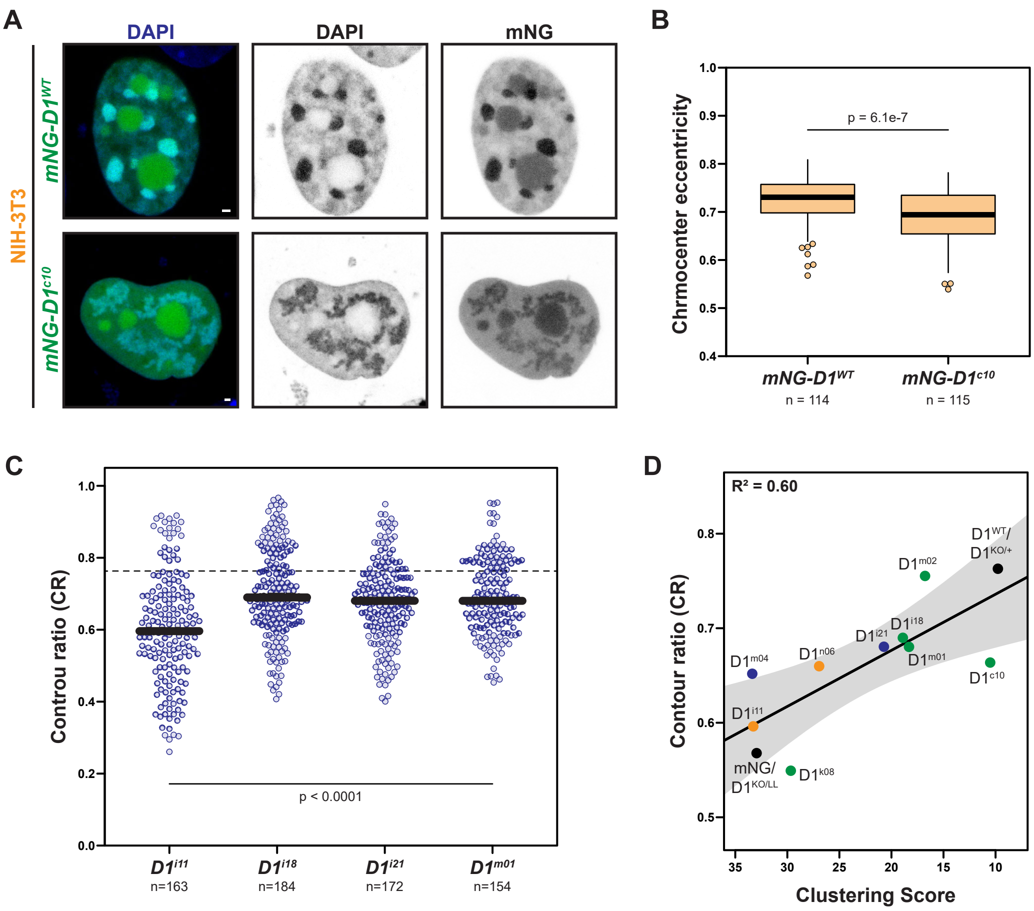


**Figure S14. Chromocenter formation is positively correlated with nuclear mechanostability.** (A) NIH-3T3 mouse fibroblast nuclei expressing $D1^{WT}/D1^{c10}$. Scale bars: 1μm. (B) Quantification of chromocenter shape in NIH-3T3 nuclei in terms of eccentricity – ratio between minor axis and major axis. (C) CR of adult spermatogonia expressing the indicated D1 variant proteins. p values are from Mann Whitney U tests in comparison to CR values from $D1^{KO/+}$ from Fig. 1C (D) Correlation between clustering scores of D1variants in mouse cells and CR measured in *Drosophila* spermatogonia. The shaded area indicates the 95% confidence intervals of the linear fit.
